# Heterogeneous Nuclear Ribonucleoprotein C is an Indispensable Target in Acute Myeloid Leukemia

**DOI:** 10.1101/2023.10.02.559746

**Authors:** Vindhya Vijay, Amy Meacham, Lauren M. Katzell, Aaron J. Winer, Jesse Terrell, Vincent L. Archibald, Lauren T. Vaughn, Leylah Drusbosky, Emily Thach, Greg Welmaker, Marcello Giulianotti, Shaun Brothers, Eleonore Beurel, Jodi L. Bubenik, Maurice S. Swanson, Alberto Riva, Abdelrahman H. Elsayed, Jatinder K. Lamba, Christopher R. Cogle

**Author notes:** Authors contributed equally (Lauren M. Katzell, Aaron J. Winer, Jesse Terrell). Authors contributed equally (Vincent L. Archibald, Lauren T. Vaughn). Correspondence to: Christopher R. Cogle, M.D. 1600 SW Archer Road Box 100278 Gainesville, FL 32610-0278.

## Abstract

Refractory disease is the greatest challenge in acute myeloid leukaemia (AML). Previously, we found vascular-associated AML cells as a source for refractory disease. In this study we use a bioassay of AML cells co-cultured on bone marrow-derived endothelial cells (BMECs) to screen 31 million compounds within a mixture-based synthetic combinatorial library followed by deconvolution leading to the identification of a novel polyamine sulfonamide that is selectively toxic to AML. Using three distinct proteomics methods we identified heterogeneous nuclear ribonucleoprotein C (HNRNPC) as the compound target.

Depleting HNRNPC reduces AML cell proliferation, viability, colony formation and fails to establish AML in mice. In contrast, no changes in cell proliferation, survival, or colony outgrowth were observed when depleting HNRNPC in normal hematopoietic or stromal cells. Clinical data show that lower HNRNPC expression independently associates with prolonged survival times in 554 adult and pediatric AML patients. We show that HNRNPC regulates DNA replication initiation and the Myc transcriptional program. Using differential isoform expression and differential alternative splicing analyses, we show significant alternative splicing of MAX after HNRNPC depletion, resulting in MAX transcript isoforms that lack Myc-interacting domains, which we term non-activating MAX isoforms. These Myc-interacting domains are necessary for the obligate heterodimerization of Myc and Max and are critical for Myc transcriptional activation. We further analyzed the alternative splicing landscapes of 538 AML patients and 33 specimens from healthy individuals and show substantial alternative splicing of mRNA in AML patients despite the absence of splicing factor gene mutations. Moreover, we found significantly high expression of non-activating MAX isoforms in healthy individuals, while AML patients highly expressed fully functional MAX isoforms, suggesting that MAX splicing may serve an important functional role in AML. Together, these data show HNRNPC as an indispensable splicing repressor in AML that is necessary for supporting deregulated Myc-Max activity and suggest HNRNPC as a potential therapeutic target for AML.

## MAIN

The greatest challenge in treating acute myeloid leukaemia (AML) is refractory disease.^1^ Previously, we identified blood vessels as a sanctuary site for chemo-refractory AML.^2^ Studies by our group and others have shown that AML cells become chemoresistant via dysregulated autocrine and paracrine signaling due to their interactions with the vascular niche.^3–5^ Targets selective to AML in the vascular niche are therefore needed for therapeutic testing. To determine vulnerabilities of vascular-associated AML cells we first screened a synthetic chemical library (SCL) comprised of over 30 million tool compounds in a co-culture assay of human AML cells (KG1) that were propagated on a layer of human bone marrow-derived endothelial cells (BMECs).

Independently, the same libraries were counter-screened with CD4+ T lymphocytes. SCLs that were highly toxic to BMECs or CD4+ T lymphocytes were discarded to reduce the odds of identifying vasculotoxic or immunosuppressive agents (Fig. 1A). Eight SCLs caused AML cell death (> 50% toxicity) while sparing underlying BMECs and were less toxic to CD4^+^ T lymphocytes (< 30% cell death) (Fig. 1B). Each SCL was iteratively interrogated by deconvolution and positional scanning to define their structure activity relationship (SAR) in relation to AML cell death and BMEC disruption. In particular, SCLs 1953 and 2210 contained the highest number of compounds that caused AML cell death while sparing underlying BMECs (Supplemental Table 1), which informed the synthesis of subsequent SCLs 2454, 2470, and 2476 (Supplemental Table 2).

**Figure 1.**
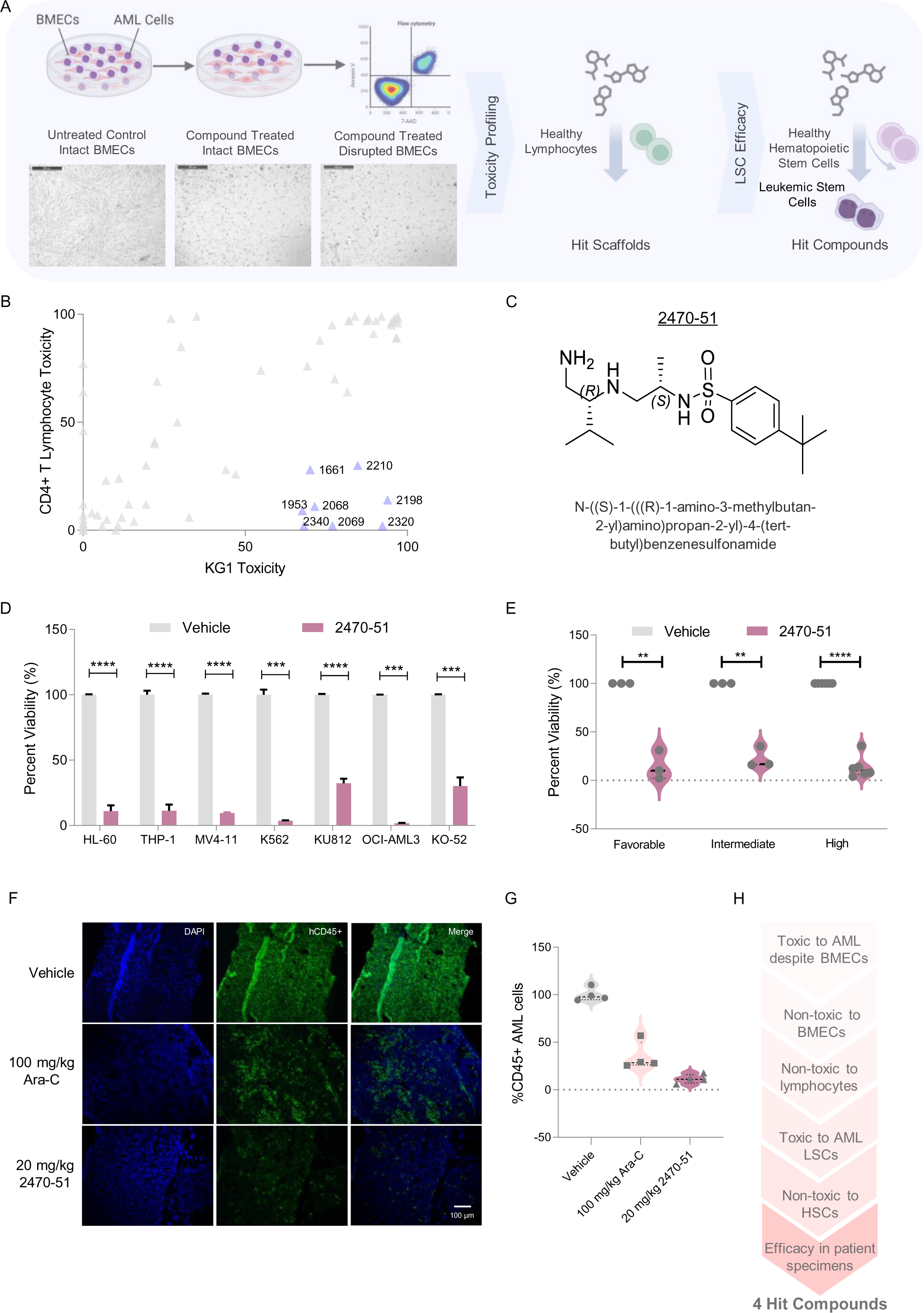
High-throughput chemical-phenotypic screening using AML-endothelial cell co-culture assay identifies anti-leukemic compounds selectively toxic to AML cells in the vascular niche. A) Schematic of drug screen and validation studies. Human AML cell line (KG-1) co-cultured on normal bone-marrow derived endothelial cells (BMECs) was used as an *in vitro* model for phenotypic screening of novel small molecule combinatorial libraries from Torrey Pines Institute of Molecular Studies (TPIMS). The treated cells were analyzed for viability using anti-Annexin V and 7-AAD using flow cytometry. Compounds that did not affect the morphology and viability of the BMECs were selected (middle micrograph) for further examination. B) Relative toxicity of screened scaffolds to CD4+ T cells and AML cell line KG-1. Compounds in the right lower quadrant were selected for further examination. C) Structure and chemical formula of hit compound 2470-51 identified from the small molecule screening. AML cell lines were treated with 2470-51 at 140 µM for 24 hours and then assessed for viability using Annexin V and 7-AAD staining followed by flow cytometry. Percentage viability of AML cell lines after 2470-51 treatment. n=3 independent experiments, error bars means ± s.e.m. *P<0.05, **P<0.01, ***P<0.001 and ****P<0.0001, Unpaired t-test. D) Percentage viability of AML patient specimens belonging to favorable, intermediate and adverse risk groups defined by 2017 ELN AML risk stratification after 2470-51 treatment. n=3, n=3 and n=6 respectively. Error bars means ± s.e.m. *P<0.05, **P<0.01, ***P<0.001 and ****P<0.0001, Unpaired t-test. F) Representative IF images of bone marrow sections from mice belonging to the three treatment groups; Saline, Cytarabine (Ara-C) at 100 mg/kg or 2470-51 at 20 mg/kg, treated intra-peritoneally TIW for 2 weeks imaged using fluorescence microscopy. Blue= DAPI nuclear staining, Green= huCD45+ AML cells and merge panel represents both DAPI and huCD45+ staining. IF images were merged using ImageJ software. G) Percentage of green huCD45+ AML cells analyzed using flow cytometric analysis of bone marrow, liver and spleens, n=4 mice per treatment group. Error bars, means ± s.e.m. *P<0.05, **P<0.01, ***P<0.001 and ****P<0.0001, Unpaired t-test. H) Screening strategy adopted to identify top hit compounds.

Individual tool compounds with high toxicity to AML cells and no disruption to BMECs were then tested for their toxicity to AML stem cells (CD34^+^CD38^-^CD123^+^) (N=6) (Supplemental Fig. 1A) and normal CD34^+^ hematopoietic stem/progenitor cells (HSPCs) (N=4). The tool compound TPI2470-51, a polyamine sulfonamide (Fig. 1C), showed the highest toxicity in AML cells and AML stem cells, no disruption to BMECs, and the greatest difference in toxicity between leukaemic and normal HSPCs (Supplemental Fig. 1B and Supplemental Table 2). Additionally, the TPI2470-51 compound was toxic in seven human AML cells lines representing various sub-types of AML (Fig. 1D). The compound was also toxic to primary AML specimens (N=12) representing all ELN risk categories (Fig. 1E), and non-toxic to normal adult bone marrow cells from healthy adult donors (N=2) (Supplemental Fig. 1C).

To assess the efficacy of the compound *in vivo*, NSG mice were transplanted with an M4 subtype human AML leukapheresis specimen with normal karyotype, *NPM1* wild type, and *FLT3* wild type (Supplemental Table 3). After 4 weeks, all mice showed human engraftment and were randomized to two weeks of treatment with either vehicle, cytarabine at 100 mg/kg IP TIW, or TPI2470-51 at 20 mg/kg IP TIW (Supplemental Fig. 2A). Two days after the last dose, mice were euthanized, and their bone marrows were examined for human CD45^+^ AML cells. We observed that mice treated with the tool compound showed significantly greater regression of AML compared to the vehicle and cytarabine control cohorts (Figure 1F, G). In addition, we observed no external signs of distress or change in body conditions due to TP12470-51 treatment (data not shown). As a preliminary *in vivo* toxicity study, we assessed the effect of TPI2470-51 on normal mouse hematopoiesis. C57Bl/6 mice were randomized to two weeks of treatment with either vehicle (N=6), cytarabine at 100 mg/kg IP TIW (N=6), or TPI2470-51 at 20 mg/kg IP TIW (N=6) (Supplemental Fig. 2B.). Mice were then euthanized, and peripheral blood and bone marrow were examined. TPI2470-51 treatment did not affect the number of WBCs or platelets (Supplemental Fig. 2C, D); however, a significant decrease in haemoglobin was observed in both cytarabine and compound-treated cohorts indicating a possibility of anaemia. Mild increase was observed in total CD45+ cells in bone marrow, liver and spleen. However, there was no gross evidence of hemorrhage or infection in these mice. A summary of compound selection approach used in this study is shown in Figure 1H.

**Figure 2.**
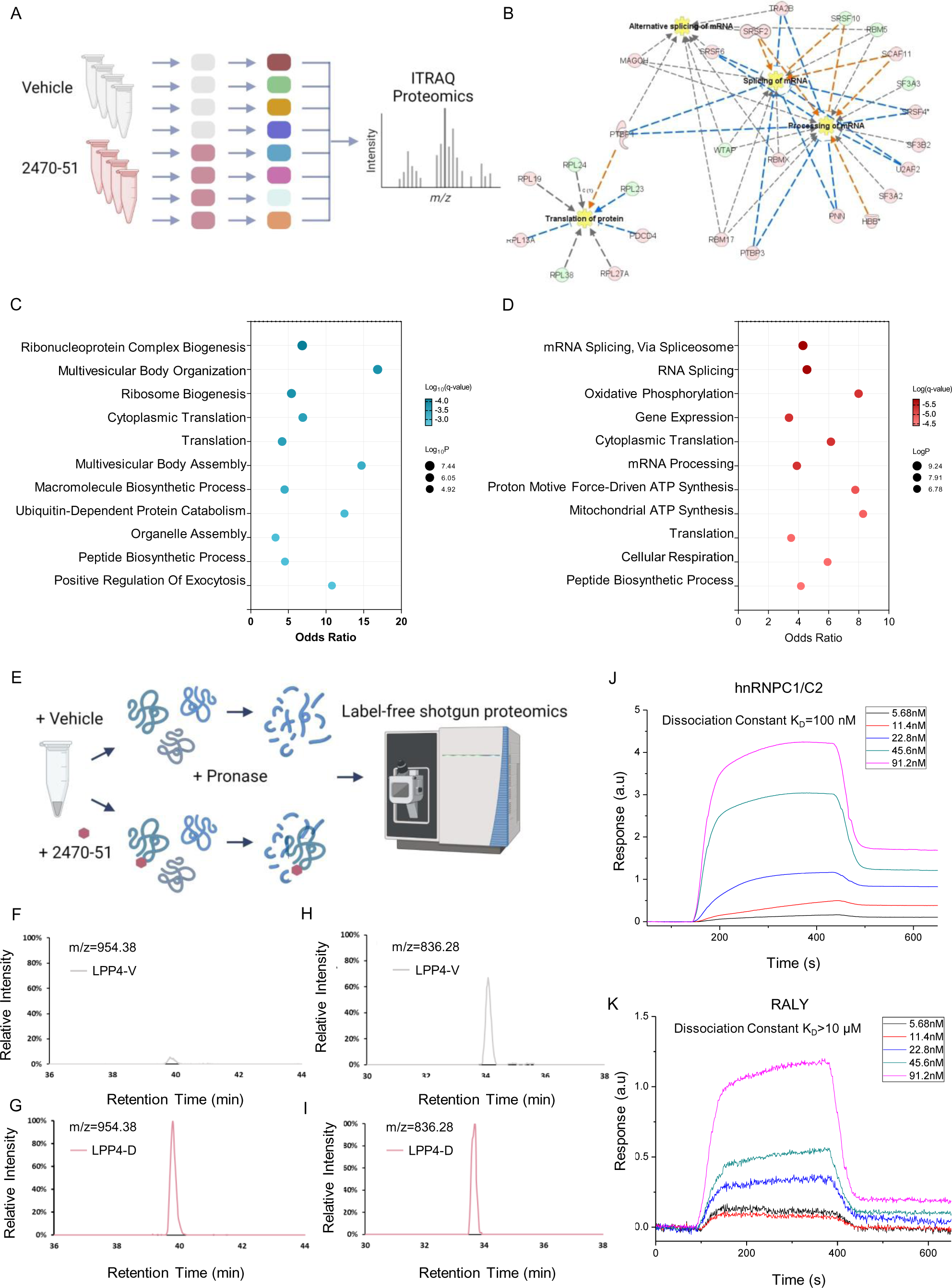
TPI2470-51 binds to HNRNPC1/C2 and alters the splicing landscape. A) Schematic of ITRAQ differential protein expression analysis performed on primary AML specimen LPP4. Each sample was treated with TPI2470-51 for 24 hours, after which proteins were isolated, labelled with isobaric tags for peptide identification and quantified using LC-MS/MS. B) Protein network analysis to identify most significantly dysregulated protein-protein networks after 24 hours of TPI2470-51 treatment as analyzed and generated using Ingenuity Pathway Analysis (IPA). The top 4 most significantly dysregulated protein networks were overlapped to generate the combined network. C-D) Gene Ontology (GO) of differentially expressed proteins after 24 hours of TPI2470-51 treatment. E) Schematic of modified DARTS approach to identify directly binding targets of TPI2470-51. LPP4 cell lysates were incubated with 500 µM 2470-51 or vehicle control for 30 minutes, followed by pronase digestion, trypsinization and analyzed using label-free shotgun proteomic method. F-G) The normalized extracted ion intensity of m/z=954.38 in 2470-51 treated (LPP4-D) and vehicle treated (LPP4-V) with the peak area of 2.78e^9^ and 1.63e^8^, respectively. The triply charged peptide at m/z 954 represents polyamine sulfonamide-associated C-terminal peptide (DDEKEAEEGEDDRDSANGEDDS) derived from trypsin digestion of a HNRNPC. H-I) The normalized extracted ion intensity of m/z=836.28 in LPP4-D and LPP4-V with the peak area of 5.18e^7^ and 3.25e^7^, respectively. The triply charged peptide at m/z 836.28 represents a C-terminal peptide (DDEKEAEEGEDDRDSANGEDDS, indicating no drug association) derived from trypsin digestion of HNRNPC. The extracted ion chromatograms were normalized against the total spectral counts of the LPP4-D and LPP4-V samples, respectively, to compensate injection variation of the two samples. J-K) Surface plasmon resonance imaging (SPRi) to identify binding affinity between immobilized recombinant human protein and 2470-51. Real-time binding signals were recorded and analyzed by Data Analysis Module (DAM). Kinetic analysis was performed using BIAevaluation 4.1 software. Association rate constant Ka, dissociation rate constant Kd and equilibrium dissociation constant KD values were calculated. J) SPRi detection of binding affinity between HNRNPC and 2470-51. K) SPRi detection of binding affinity between RALY and 2470-51

To identify the targets of the tool compound, we first analyzed the differential protein expression in AML cells (primary specimen LPP4, Supplemental Table 4) treated with vehicle control versus TPI2470-51 and then analyzed at 24 hours by multiplexed isobaric labeling and tandem mass spectrometry (ITRAQ)^6^ (Fig. 2A). Using LC-MS/MS analysis, we identified 108,925 expressed peptides corresponding to 3,079 proteins. Of these proteins, 2,014 proteins were differentially expressed between vehicle versus TPI2470-51-treated samples (Supplemental Fig. 2A). Protein-protein interaction (PPI) analysis of the differentially expressed proteins identified splicing of mRNA, alternative splicing of mRNA, processing of mRNA, and translation as the top four most significantly dysregulated network of proteins (Fig. 2B). Pathway and process enrichment analysis indicated that the top dysregulated pathways were metabolism of RNA, RNA splicing, ribonucleoprotein complex biogenesis, and translation (Fig. 2C, D Supplemental Table 5).

To pinpoint binding targets, we footprinted the tool compound by adapting a drug affinity responsive target stability (DARTS) method using tandem mass spectrometry for peptide analyses.^7–9^ Proteins isolated from human AML (primary specimen LPP4, Supplementary Table 3) were incubated with the tool compound or vehicle control, and then digested with pronase (Fig. 2E). Peptides were then analyzed by a label-free shotgun proteomic method (Supplemental Methods). We then identified proteins whose stability may be affected by the compound using LC-MS/MS (Supplemental Methods). This analysis led to the identification of ten proteins interacting with TPI2470-51 (Supplemental Table 6). Both enolase and heterogeneous nuclear ribonucleoprotein C (HNRNPC) showed more spectra in the compound treatment compared to vehicle control, suggesting compound-binding enhanced the stability of the protein (Supplemental Table 7). Based on the differential protein expression data showing perturbations in RNA splicing, we chose to focus our examination on HNRNPC, an RNA-binding protein involved in modulating mRNA stability, splicing and translation.^10–15^ Our proteomic analyses revealed a MS/MS spectrum for the C-terminal peptide of HNRNPC. On scanning and comparing the C-terminal peptide of vehicle versus TPI2470-51-treated specimens, we identified the peptide DDEKEAEEGEDDRDSANGEDDS (m/z=954.38) to be selectively changed by the compound (Fig. 2F, G). However, an unrelated peptide at m/z=836.28 remained unchanged between vehicle and TPI2470-51 treated specimens (Fig. 2H, I). As validation of these findings, surface plasmon resonance imaging (SPRi) (Supplemental Methods) showed a lower equilibrium dissociation constant (K_D_) for the interaction between the tool compound and HNRNPC of 1.05 x 10^-7^M (∼100 nM) (Fig. 2J), in contrast to RALY, a ribonucleoprotein with high sequence homology (∼69%) to HNRNPC^16, 17^ (Fig. 2K) and an equally abundant ribonucleoprotein, hnRNP A2/B1 (Supplemental Fig. 3A), which showed K_D_ values that were 50-to 150-fold higher, respectively (Supplemental Table 8). These data led us to hypothesize that HNRNPC is a molecular and functional target for the anti-leukemic compound TPI2470-51.

**Figure 3.**
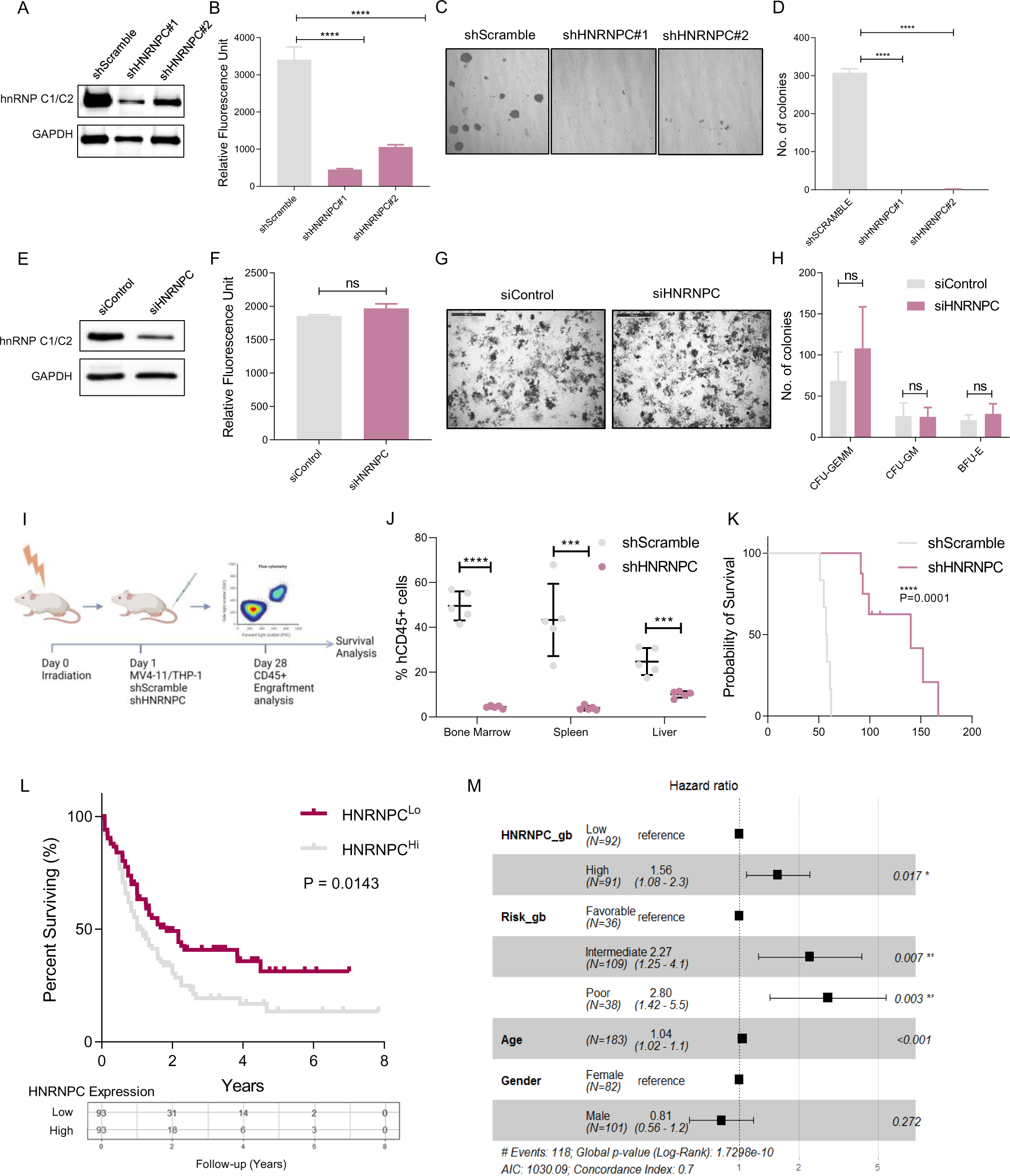
HNRNPC depletion reduces AML growth *in vitro* and *in vivo*. A) AML cell line K562 was transduced with lentiviruses expressing either a scrambled control (shSCRAMBLE) or two independent shRNA targeting HNRNPC (shHNRNPC#1 and shHNRNPC#2) for 48 hours, followed by puromycin selection for 3-7 days. Puromycin resistant cells were immunoblotted for HNRNPC and Gapdh (loading control), day 5 after transduction. B) AML cell metabolic activity using Cell titer blue assay as a surrogate for cell proliferation 7d post-transduction. Y-axis represents relative fluorescent units (RFU). n=3 independent experiments, error bars mean ± s.e.m. *P<0.05, **P<0.01, ***P<0.001 and ****P<0.0001, Unpaired t-test. C) Representative images of colony forming unit (CFU) assay of K562 cells 14d post-selection in a methylcellulose based medium. D) Number of colonies (>50 cells) counted 14d after plating in methylcellulose medium. Error bars means ± s.e.m. *P<0.05, **P<0.01, ***P<0.001 and ****P<0.0001, Unpaired t-test. E) Mononuclear cells isolated from human UCB specimens were transduced with siRNA targeting a scrambled control (siControl) or HNRNPC (siHNRNPC) for 72 hours. Cells were immunoblotted for HNRNPC and Gapdh (loading control). F) UCB cell metabolic activity using Cell titer blue assay as a surrogate for cell proliferation. Y-axis represents relative fluorescent units (RFU). n=3 independent experiments, error bars means ± s.e.m. *P<0.05, **P<0.01, ***P<0.001 and ****P<0.0001, Unpaired t-test. G) Representative images of colony forming unit (CFU) assay of UCB cells 14d after plating in a methylcellulose based medium. H) Number of colonies (>50 cells) counted for CFU-granulocyte, erythrocyte, monocytes and macrophages (CFU-GEMM), CFU-granulocyte and monocytes (CFU-GM), and blast forming unit-erythrocytes (BFU-E). Error bars means ± s.e.m. *P<0.05, **P<0.01, ***P<0.001 and ****P<0.0001, Unpaired t-test. I) MV4-11 cells were transduced with lentiviruses expressing either a scrambled control (shScramble) or shRNA targeting HNRNPC (shHNRNPC#1 & #2) for 48 hours. Puromycin resistant cells 5d post-transduction were transplanted into irradiated NSG mice. 6 weeks post transplantation, mice were euthanized and organs were harvested for MV4-11 engraftment analysis. J) Quantification of hCD45+ cells in bone marrow, liver and spleen analyzed using flow cytometry. n=5 mice per cohort, error bars means ± s.e.m. *P<0.05, **P<0.01, ***P<0.001 and ****P<0.0001, Unpaired t-test. K) THP-1 cells were transduced with lentiviruses expressing either a scrambled control (shScramble) or shRNA targeting HNRNPC (shHNRNPC#1 & #2) for 48 hours. Puromycin resistant cells 5d post-transduction were transplanted into irradiated NSG mice and assessed for overall survival. Kaplan-Meier estimate for overall survival n=6, error bars means ± s.e.m. *P<0.05, **P<0.01, ***P<0.001 and ****P<0.0001, Unpaired t-test. L) Survival probabilities of TCGA-AML cohort based on HNRNPC expression. M) OS probabilities of patients based on HNRNPC expression. Forest plot indicating the hazard ratios and P-values for each variable used in the multi-variate survival analysis.

To interrogate the function of HNRNPC in AML, we depleted HNRNPC in AML cell lines using two independent shRNA constructs that knock-down HNRNPC (HNRNPC-kd). Depleting HNRNPC led to significant decreases in proliferation and viability in human AML cell lines K562 and THP-1 (Figure 3A, B and Supplementary Fig. 4A, B) Furthermore, HNRNPC depletion resulted in significantly decreased AML colony forming units (Figure 3C, D and Supplementary Fig. 4C, D). In contrast, HNRNPC depletion in normal hematopoietic cells from umbilical cord blood specimens, primary BMECs and fibroblastic stromal cells showed no changes in proliferation, viability or colony outgrowth (Figure 4E-H and Supplemental Fig. 4E-I). Together, these data indicate that HNRNPC is indispensable in AML cells, yet dispensable for normal cell survival, which agrees with a previous reports showing the dispensability of HNRNPC in mouse embryonic stem cells barring effects on neurodevelopment.^18, 19^

**Figure 4.**
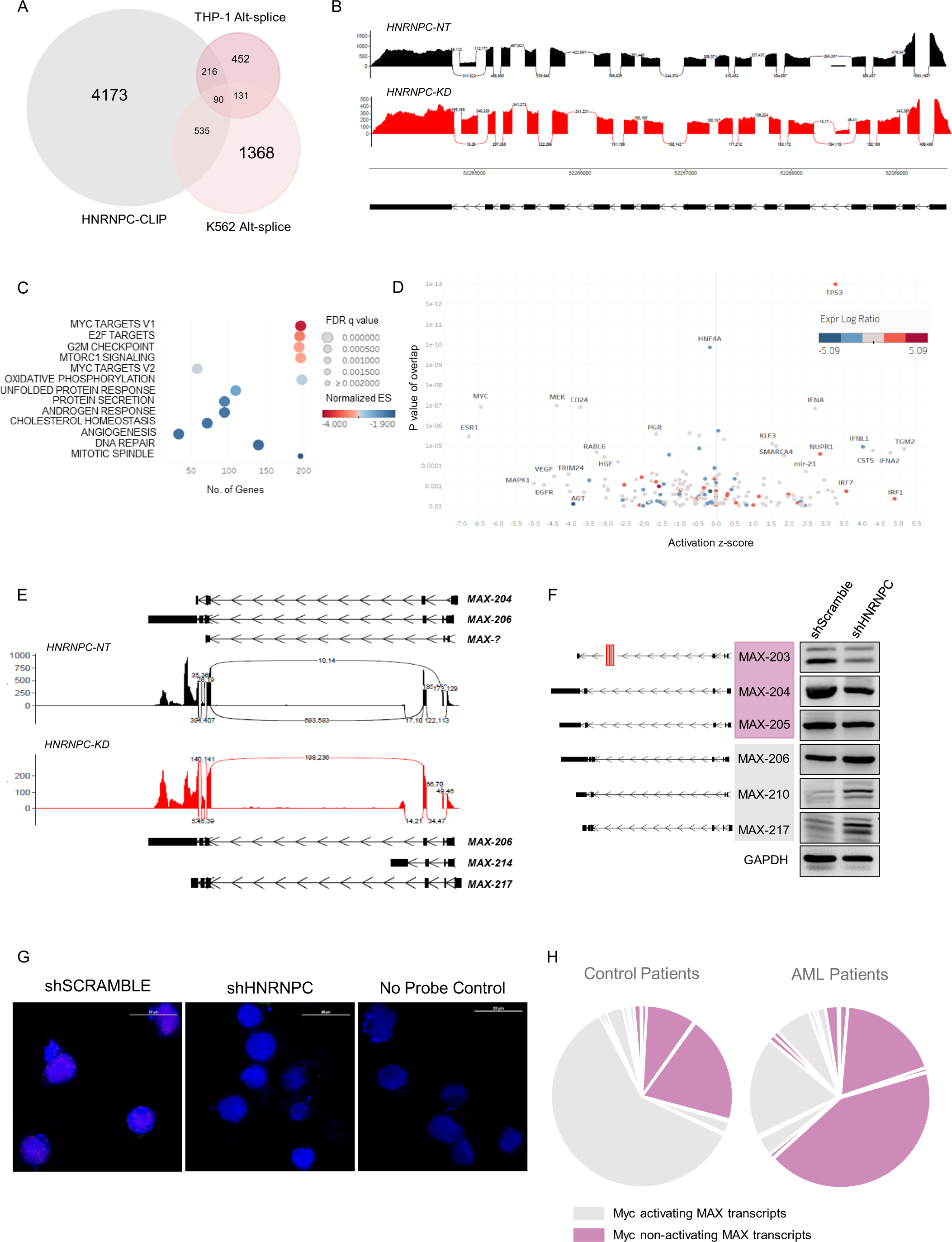
HNRNPC1/2 regulates MYC transcriptional activity via mis-splicing of MAX mRNA. A) Comparison of THP-1, K-562 alternative splicing events after HNRNPC depletion and HNRNPC CLIP-Seq data. Venn diagram indicates common events that are mediated by HNRNPC. B) Effect of HNRNPC depletion on the splicing pattern of MCM3. Sashimi plots represent the splicing pattern of MCM3 in THP-1 cells after HNRNPC depletion. C) Gene-set enrichment analysis (GSEA) of genes differentially expressed after HNRNPC depletion. D) Upstream regulator analysis of differentially expressed genes after HNRNPC depletion in THP-1 cells using Ingenuity® Pathway Analysis (IPA). The activation z-score infers the predicted activation state of upstream transcription factors based on the overlap between experimental trends and Ingenuity® knowledge base. P-Value of overlap represents the significance of overlap between the two datasets. E) Effect of HNRNPC depletion on splicing patterns of MAX. Sashimi plot representing the splicing pattern of MAX in HNRNPC-NT (Black) and HNRNPC-kd (Red) cells. F) RT-PCR validation of MYC-activating (pink) and MYC-inactivating (grey) MAX isoforms. The black lines represent the transcript map for each MAX isoform. The red double line indicates that the intron has been segmented to fit within space restrictions. G) Proximity ligation assay assessing Myc and Max interactions in HNRNPC*-*NT and HNRNPC*-*Kd cells. The negative control is AML cells that were incubated with only anti-Myc primary antibody. H) Pie chart representing proportional distribution of activating and non-activating MAX isoforms in healthy (control) patients and AML patients. Each section represents an individual MAX transcript.

We next evaluated the functional importance of HNRNPC in AML by comparing *in vivo* engraftment of AML HNRNPC-kd versus HNRNPC-NT cells. Human AML cells (MV4-11) with HNRNPC depletion failed to engraft in hematopoietic tissues (i.e., bone marrow, spleen, liver) of sub-lethally irradiated mice compared to AML cells with wild-type (WT) HNRNPC expression (Figure 3I, J and Supplemental Fig. 4J). Additionally, mice transplanted with HNRNPC*-*kd cells showed significantly longer survival in comparison to mice transplanted with HNRNPC*-*NT cells (Figure 3K). Together, these *in vivo* experiments show the functional necessity of HNRNPC in AML.

To examine the extent by which HNRNPC expression influences AML patient prognosis, we studied the survival times of a total of 554 patients in three AML cohorts (TARGET, AML-02, TCGA^20^) with respect to their AML HNRNPC expression and in context to other known prognostic variables like age, gender and genetic risk stratification. In both adult AML (TCGA) (Figure 3L) cohorts and pediatric (TARGET, AML-02) (Supplemental Fig. 5A-D and Supplemental Table 9), lower HNRNPC expression in AML cells was associated with prolonged survival and was an independent predictor of survival (Figure 3M).

To investigate the molecular mechanisms controlled by HNRNPC in AML cells, we first compared gene expression changes between HNRNPC-NT vs. HNRNPC-kd THP1 cells (Supplemental Table 10). In addition, we also analyzed publicly available RNA-Seq dataset (ENCODE) for K562 cells with HNRNPC depletion. The top molecular pathways with dysregulation were processes involved in DNA replication initiation and cell cycle transition (Supplemental Fig. 6A). To identify aberrant splicing events contributing to dysregulation of gene expression, we performed alternative splicing analysis on both THP1 and K562 datasets. Using HNRNPC iCLIP data^21^ and comparing with significantly alternatively spliced transcripts, we identified HNRNPC-associated splicing events that were specifically altered by HNRNPC depletion (Figure 4A and Supplemental Table 11). Depleting HNRNPC in THP-1 cells led to significant mis-splicing of DNA-replication licensing factor *MCM3*. We observed that HNRNPC depletion resulted in aberrant inclusion of a region within Intron 16 (Figure 4B), which has previously been reported to contain Alu elements that are highly susceptible to degradation via nonsense-mediated decay (NMD).^22^ As expected, we observed that HNRNPC depletion led to inclusion of intron 16 in MCM3 mRNA (Supplemental Fig. 6 B, C) which led to a significant decrease in MCM3 protein expression (Supplemental Fig. 6D).

Gene set enrichment analysis (GSEA) showed that the most significantly dysregulated genes were highly enriched in gene sets containing MYC targets, E2F targets, G2M targets, and MTORC signaling (Figure 4C). To identify transcriptional regulators resulting in this pattern of gene expression, IPA upstream regulator analysis was performed, and we recognized inactivation of MYC, ESR1, MEK and CD24 adhesion molecule and activation of TP53 (Figure 4D Supplemental Table 12). Since GSEA identified enrichment of MYC transcriptional targets and IPA recognized MYC as an inactivated upstream transcriptional regulator, we hypothesized that HNRNPC depletion critically dysregulates the MYC transcriptional program in AML cells. Interrogation of mRNA expression confirmed decreased expression of MYC targets in AML-HNRNPC-kd cells compared to controls (Supplemental Fig. 7A). However, neither *MYC* mRNA expression nor Myc splicing changed when HNRNPC was downregulated (Supplemental Fig. 7B, C). Myc-induced transcription requires successful heterodimerization between Myc and its obligate partner Max to bind targets with E-box consensus sequence.^23^ Myc and Max proteins interact with each other using their bHLH and LZ domains to form a coiled-coil structure capable of binding DNA. Prior studies have established that lack of Myc-Max heterodimerization significantly disrupts Myc-induced transcription^24, 25^, which is a key mechanism for oncogenic signaling in AML.^26–28^ Thus, we examined Myc’s obligate partner, Max, and found significant decrease in protein expression (Supplemental Fig. 7D) in HNRNPC-kd cells. RT-PCR analysis showed that depleting HNRNPC resulted in an increase of MAX transcripts that lacked bHLH domains, whereas isoforms that contain bHLH domain where significantly decreased (Figure 4E). Our results pointed towards the possibility of an isoform switch, where HNRNPC depletion favored the production of Myc-inactivating MAX transcripts and decrease of Myc-activating MAX transcripts.

MAX transcripts bind to HNRNPC, as evidenced by CLIPseq^21^. HNRNPC downregulation in AML cells not only decreases MAX expression but also produces alternatively spliced (AS) transcripts fated for truncated structures or nonsense mediated decay (NMD) (Figure 4F). We hypothesized that Max isoforms resulting from these mis-spliced MAX transcripts after HNRNPC depletion were incapable of heterodimerizing with Myc in AML cells. To interrogate this, proximity ligation assay (PLA) showed significantly fewer functional Myc-Max interactions when HNRNPC was downregulated in AML cells (Figure 4G and Supplemental Fig. 7E). These results confirm that aberrant MAX splicing disrupts Myc-Max heterodimerization, consequently affecting the ability of Myc to bind to DNA. Taken together, our results validate that Myc transcriptional program is inactivated upon HNRNPC depletion due to disruption of Myc-Max heterodimerization caused by mis-splicing of MAX mRNA.

To determine clinical relevance, we examined the Beat AML dataset of 538 AML patients.^29^ We observed that *MYC* and MAX gene expression were not significantly different between AML patients and healthy controls (Supplemental Fig. 8A, B). Next, we assessed the splicing patterns of this cohort and identified significant mis-splicing in AML patients in comparison to the healthy controls (Supplemental Fig. 8C). We observed that the majority of the mis-spliced transcripts were enriched in pathways involved in DNA replication, cell cycle regulation and translation (Supplemental Fig. 8D, E). While *MYC* splicing was not significantly altered between AML patients and healthy controls (Supplemental Fig. 8F, G), we found that MAX splicing was significantly altered. Healthy control patients expressed a higher proportion of MAX transcripts incapable of interacting with Myc compared to AML patients that expressed a higher proportion of activating MAX transcripts (Figure 4H and Supplemental Table 13). Our results indicate that the splicing program in AML is inherently altered to enable production of activating MAX spliced isoforms that facilitate Myc-Max dimerization. This abnormal alteration in splicing pattern, at least partially, explains the hyperactive Myc activity, despite no apparent gene expression changes. Furthermore, we were able to draw a correlation between our in vitro findings and AML clinical data. With HNRNPC depletion, we observed that MAX splicing favored increased production of non-activating isoforms and a decrease in activating MAX isoforms. Now, we see that AML aberrantly expresses activating MAX isoforms which upon HNRNPC depletion should alter the splicing landscape of AML to express non-activating isoforms, as occurring in healthy patients. While AML cells will not tolerate this isoform switching leading to decrease in critically required Myc transcriptional program, we suspect that normal cells, primarily expressing non-activating isoforms will be unperturbed by the splicing changes, potentially explaining the selective dependence of AML on HNRNPC expression.

In this study, we start with a phenotypic screen for compounds capable of eliminating AML cells embedded in the chemo-protective vascular niche, identify a splicing repressor necessary for AML cell proliferation and survival, define a mechanistic role for HNRNPC in leukaemia, and exhibit HNRNPC depletion as a new, and potentially therapeutic strategy for treating AML. The findings presented here provide considerable new insight into the importance of HNRNPC in AML, in contrast to its dispensable role in normal hematopoietic and stromal cells. Together, these data define HNRNPC as a new candidate target for the treatment of AML and offer a pharmacological agent for engaging the target.

## ACKNOWLEDGEMENTS

We thank all the patients who donated specimens and data to The Malignant Hematology Bank (UF IRB#201501063). Funding for this project was provided in part by the Gatorade Foundation administered by the University of Florida Department of Medicine. CRC received funding from a Scholar in Clinical Research Award from the Leukemia & Lymphoma Society, the Spanier Foundation, the Harry T. Mangurian Foundation, a Pierre Chagnon Professorship, and the Oxnard Family Foundaton. We thank Dr. Richard Houghten at Torrey Pines Institute for Molecular Studies for his advice on library screenings. We also thank Dr. Lucas J. Boatwright at University of Florida ICBR for his support with bioinformatic analyses.

## AUTHOR CONTRIBUTIONS

V.V. led the project, performed experiments, analyzed data and wrote the manuscript. A.M., L.K., A.J.W. and J.T., L.V., V.A. and E.T. performed experiments and analyzed data. L.D. S.B. and E.B. provided experimental support. G.W. and M.G. provided the synthetical compound libraries and medicinal chemistry guidance. J.L.B., M.S.S., A.R., A.H.E. and J.L. performed bioinformatic analyses and edited the manuscript. C.R.C. provided project oversight for experimental design, data management, data analysis and wrote the manuscript.

## METHODS

### Cell Culture

Human AML cell lines KG-1 and HL-60 were cultured in IMDM supplemented with 20% fetal bovine serum (FBS) and antibiotics. MV4-11 and K562 cell lines were cultured in IMDM supplemented with 10% FBS and antibiotics. THP-1 cells were cultured in IMDM supplemented with 10% FBS, antibiotics and 0.5 mM b-mercaptoethanol (BME). K0-52, OCI-AML3, KU812 cell lines and human lung cancer cell line A549 were cultured in RPMI supplemented with 10% FBS and antibiotics. Finally, human triple-negative breast cancer cell line MDA-MB-231-LM2 and HEK293T were cultured in DMEM supplemented with 10% FBS and antibiotics.

Primary AML specimens included in the study were bone marrow or peripheral blood from patients with AML. The University of Florida Health Center Institutional Review Board (IRB) approved use of these human samples under protocols #201501063 (The Malignant Hematology Bank) and #201600248 (Laboratory Studies of Normal and Malignant Cells). Primary AML cells from leukemic bone marrow specimens, mononuclear cells from healthy bone marrow specimens (Lonza) or cord blood cells from human umbilical cord blood specimens (Life South) were cultured in Stemspan supplemented with necessary cytokines. Primary BMECs derived from healthy bone marrow specimens were cultured in EBM-2 media (Lonza) supplemented with necessary cytokines (Lonza).

### *In Vitro* Chemical-Phenotypic Screening Using AML-BMEC Co-Culture Model

Briefly, BMECs were seeded in a 24-well plate for 24-48 hours. At 70% confluency, media was aspirated and 300,000 AML cells were seeded in each well. After 2-4 hours, desired compounds were added to the cells and allowed to incubate for 24 hours. The cells were then collected and analyzed using flow cytometry to quantify AML cell viability. Phenotypic screening of approximately 31 million compounds were performed using the AML-BMEC co-culture assay. Compound libraries dissolved in 20% Dimethyl Formamide (DMF) were diluted in IMDM or RPMI to make a final concentration of 50 µg/mL. The cells were treated with each scaffold or vehicle (20% DMF) in triplicates for 24 hours. At the end of 24 hours, AML cells were harvested and stained with human CD45-V450, Annexin V-FITC and 7-AAD antibodies (1:50 uL PBS, BD Biosciences) for analysis using flow cytometry. For analyzing the stem and progenitor cell population, additional antibodies human CD34-PE, CD38-FITC and CD123-BV421 (1:50 uL PBS, BD Biosciences) were added to the panel for flow cytometric analysis. The compound libraries were synthesized using a positional scanning deconvolution method. ^28^ Individual compounds were synthesized after identification of hit scaffolds with desired characteristics. Hit compounds were rank-ordered based on AML and normal cell toxicity.

### Purification of Individual Compounds

All purification were carried out using a Shimadzu Semi-preparative HPLC. The HPLC system consisting of LC-6AD binary pumps, a DGU-20A 3R degassing unit, a CBM-20A communication bus module, a SIL-10AP auto sampler, and a FRC-10A fraction collector. All chromatographic peak detections were performed by using a SPD-20A diode array detector set to detect 214nm absorbance. All chromatographic separations were performed using a Phenomenex Luna C18 preparative column (5µm, 150 x 21.20mm i.d.). The inlet of the column was protected by a Phenomenex C18 Prep security guard cartridge (15 x 21.2mm i.d.).

All samples were reconstituted in 2000/3000uL of a 50/50 mixture of HPLC grade or higher acetonitrile/water mixture obtained through Sigma Aldrich. The samples were then filtered through a Spartan 30, 0.45 µm syringe filter before injecting onto the HPLC for separation. The mobile phases consisted of deionized water and HPLC grade acetonitrile with 0.1% LCMS grade formic acid obtained from Sigma Aldrich and Fisher Scientific.

The initial setting for the HPLC was 95% water. The gradient was linearly increased to 20% acetonitrile (v/v) over 6 minutes. The gradient was again linearly increased to 60% acetonitrile (v/v) over 39 minutes. The gradient was then linearly increased to 95% acetonitrile (v/v) over 3 minutes and then held for an additional 4 minutes. Finally, the gradient was linearly decreased to 5% acetonitrile (v/v) over 1 minute and held until stop. The total run time was equal to 58 minutes. The total flow rate used was 12mL per minute and peak divide time was set at 0.75 minutes. A slope of 1,000,000uV/sec was set for this unit along with a level of 1,004,500uV, a slope override was also set for 5,510uV. Sample volume injected were based upon crude material yields with either 2000uL or 3000uL being injected 1x.

### General Synthesis of Positional Scanning Library 2210 and Individual Compounds 2470

Library 2210 and all the individual polyamine sulfonamides reported within were synthesized following the same general scheme (as previously reported^29^). The solid phase synthesis was performed using the “tea bag” methodology.^30, 31^ Initially, 100 mg of *p*-methylbenzdrylamine (MBHA resin (1.1 mmol/g, 100-200 mesh) was sealed in a mesh “tea-bag,” neutralized with 5% diisopropylethylamine (DIEA) in dichloromethane (DCM) and subsequently swelled with additional DCM washes. Boc-amino acids (6 equivalent) were coupled in DMF (0.1M DMF) for 120 min in the presence of diisopropylcarbodiimide (DIC, 6 equiv) and 1-hydroxybenzotriazole hydrate (HOBt, 6 equiv). Boc-amino acids (6 equivalent) were coupled in DMF (0.1M DMF) for 120 min in the presence of diisopropylcarbodiimide (DIC, 6 equiv) and 1-hydroxybenzotriazole hydrate (HOBt, 6 equiv). The Boc protecting group was removed with 55% TFA/DCM for 30 min and subsequently neutralized with 5% DIEA/DCM (3x) (1, Supplementary Fig. 1D)). Boc-amino acids were coupled utilizing standard coupling procedures (6 equiv) with DIC (6 equiv) and HOBt (6 equiv) in DMF (0.1M) for 120 min. The Boc protecting group was removed with 55% TFA/DCM for 30 min and subsequently neutralized with 5% DIEA/DCM (3x) (2, Supplementary Fig. 1D). The sulfonamide was formed using Sulfonyl chlorides (10 equiv) and DIEA (10 equiv) in 0.1 M DCM for 120 min. The bags were washed with DCM (3x) (3, Supplementary Fig. 1D). All coupling reactions were monitored for completion by the ninhydrin test. The reduction was performed in a 4000 mL Wilmad LabGlass vessel under nitrogen. A borane in 1.0 M tetrahydrofuran complex solution was used in 40-fold excess for each amide bond. The vessel was heated to 65 °C and maintained at temperature for 72 h. The solution was then discarded, and the bags were washed with THF and methanol. Once completely dry, the bags were treated overnight with piperidine at 65 °C and washed several times with methanol, DMF, and DCM (4, Supplementary Fig. 1D). Before proceeding, completion of reduction was monitored by a control cleavage and analyzed by LCMS. The resin was cleaved with HF in the presence of anisole in an ice bath at 0 °C for 7 h (5, Supplementary Fig. 1D). The positional scanning library incorporates both individual and mixtures of amino acids (R1 and R2) and sulfonyl chlorides (R3). The synthetic technique facilitates the generation of information regarding the likely activity of individual compounds from the screening of the library. Equimolar isokinetic ratios have been previously determined and calculated for each of the amino acids and sulfonyl chlorides utilized for the respective mixtures. The polyamine sulfonamide library 2210 has a total diversity of 17,340 compounds (34 x 34 x 15 = 17,340).

### *In vivo* efficacy studies in AML PDX models

All animal studies were performed according to approved protocols from the University of Florida Institutional Animal Care and Use Committee (IACUC). To test the efficacy of hit compound 2470-51, we used 6-8 weeks old immuno-deficient NSG (NOD.Cg-*Prkdc^scid^Il2rg^tm1Wjl^*/SzJ) mice as xenograft hosts for primary AML samples. Following sub-lethal irradiation (200 cGy), NSG mice received tail vein injections of 2 x 10^6^primary human AML cells. After 4 weeks of engraftment, the animals were randomized into three groups: vehicle-treated (negative treatment control), cytarabine-treated 100 mg/kg IP TIW (positive treatment control) and hit compound 20 mg/kg IP TIW. After two weeks of treatment, all cohorts were euthanized to analyze for human AML engraftment. Cells from bone marrow, blood, spleen and liver from each animal was examined for the presence of human CD45+ AML blasts and CD34+/CD38-AML LSCs using flow cytometry. Additionally, femur bones, liver and spleen tissues were fixed in 4% PFA, decalcified using EDTA if required and embedded in OCT for further immunofluorescence staining. The OCT cryosections were stained with human CD45-FITC and DAPI nuclear stain to visualize AML engraftment.

### Differential Protein Expression

Protein extracts from three AML specimens treated with vehicle or TPI2470-51 for 24 hours were collected and quantified as previously described (reference original iTRAQ manuscript). Proteins were then dissolved in denaturant buffer (0.1% SDS (w/v)) and dissolution buffer (0.5 M triethylammonium bicarbonate, pH 8.5) in the iTRAQ Reagents 8-plex kit (AB sciex Inc.). For each sample, a total of 100 μg of protein were reduced, alkylated, trypsin-digested, and labeled according to the manufacturer’s instructions (AB Sciex Inc.). The vehicle- and compound-treated samples were labeled with unique iTRAQ tags. Labeled peptides were desalted with C18-solid phase extraction and dissolved in strong cation exchange (SCX) solvent A (25% (v/v) acetonitrile, 10 mM ammonium formate, and 0.1% (v/v) formic acid, pH 2.8). The peptides were fractionated using an Agilent HPLC 1260 with a polysulfoethyl A column (2.1 × 100 mm, 5 µm, 300 Å; PolyLC.). Peptides were eluted with a linear gradient of 0–20% solvent B (25% (v/v) acetonitrile and 500 mM ammonium formate, pH 6.8) over 50 min followed by ramping up to 100% solvent B in 5 min. The absorbance at 280 nm was monitored and a total of 12 fractions were collected. The fractions were lyophilized and re-suspended in LC solvent A (0.1% formic acid in 97% water (v/v), 3% acetonitrile (v/v)). A hybrid quadrupole Orbitrap (Q Exactive) MS system (Thermo Fisher Scientific, Bremen, Germany) was used with high energy collision dissociation (HCD) in each MS and MS/MS cycle. The MS system was interfaced with an automated Easy-nLC 1000 system (Thermo Fisher Scientific). Each sample fraction was loaded onto an Acclaim Pepmap 100 pre-column (20 mm × 75 μm; 3 μm-C18) and separated on a PepMap RSLC analytical column (250 mm × 75 μm; 2 μm-C18) at a flow rate of 350 nl/min during a linear gradient from solvent A (0.1% formic acid (v/v)) to 30% solvent B (0.1% formic acid (v/v) and 99.9% acetonitrile (v/v)) for 95 min, to 98% solvent B for 15 min, and hold 98% solvent B for additional 30 min. Full MS scans were acquired in the Orbitrap mass analyzer over m/z 400–2000 range with resolution 70,000 at 200 m/z. The top ten most intense peaks with charge state ≥ 3 were fragmented in the HCD collision cell normalized collision energy of 28%, (the isolation window was 2 m/z).

The maximum ion injection times for the survey scan and the MS/MS scans were 250 ms respectively and the ion target values were set to 3e6 and 1e6, respectively. Selected sequenced ions were dynamically excluded for 60 sec. To reduce the probability of false peptide identification, we only included peptides at the 95% confidence interval by a ProteinPilot probability analysis, and each confident protein identification was supported by at least one unique peptide. We only used ratios with p-values ≤ 0.05, and only fold changes of > 1.5 or < 0.5 were considered as significant. Functional enrichment analysis and pathway analysis of differentially expressed proteins were performed using an online tool called Metascape. Protein-protein interaction networks were built using Ingenuity Pathway Analysis (IPA) (Qiagen).

### Differential Protein Expression Analysis

The raw MS/MS data files were processed by a thorough database searching approach considering biological modification and amino acid substitution against UNIPROT Human database (148,284 entries; downloaded on Oct. 9th, 2015) using the ProteinPilot v4.5 with the Fraglet and Taglet searches under ParagonTM algorithm. The following parameters were considered for all the searching: fixed modification of methylmethane thiosulfonate-labeled cysteine, fixed iTRAQ modification of amine groups in the N-terminus, lysine, and variable iTRAQ modifications of tyrosine. For protein quantification, only MS/MS spectra that were unique to a particular protein and where the sum of the signal-to-noise ratios for all the peak pairs > 9 were used for quantification.

### Sample preparation and electrophoresis-Compound footprinting using DARTS

After denaturation using boiling and cooling-down on ice, 30 μl of the clear supernatant was applied to a 12% acrylamide SDS page gel (BioRad, Hercules, CA) and run for 45 min at 200 V. The gels were washed three times in Milli-Q water for 5 min each time and stained overnight in 20 ml Invitrogen SimplyBlue^™^ SafeStain with gentle shaking. The excised bands from the gel were de-stained with 1ml 50 mM ammonium bicarbonate, pH 8.0/acetonitrile (1:1, v/v). Each sample was reduced with 40 mM DTT, alkylated with 100 mM of 2-chloroacetamide, and trypsin-digested. Tryptic digested peptides were desalted with C18-Ziptip (Millipore, Billerica, MA, USA).

### MS/MS analysis

The hybrid quadrupole Orbitrap (Q Exactive Plus) MS system (Thermo Fisher Scientific, Bremen, Germany) was interfaced with an automated Easy-nLC 1200 system (Thermo Fisher Scientific, Bremen, Germany).

Samples was loaded onto an Acclaim Pepmap 100 pre-column (20 mm × 75 μm; 3 μm-C18) and separated on a PepMap RSLC analytical column (250 mm × 75 μm; 2 μm-C18) at a flow rate at 250 nl/min during a linear gradient from solvent A (0.1% formic acid (v/v)) to 25% solvent B (0.1% formic acid (v/v), 19.9% water (v/v) and 80 % acetonitrile (v/v)) for 80 min, and to 100% solvent B for additional 15 min. The spectrum library was produced in the data dependent mode with survey scans acquired at a resolution of 70,000 at 200 m/z. The mass spectrometer was operated in MS/MS mode scanning from 350 to 1800 m/z. Up to the top 10 most abundant isotope patterns with charge 2 ∼ 5 from the survey scan were selected with an isolation window of 1.2 Th and fragmented by high energy collision dissociation (HCD) with normalized collision energies of 28. The maximum ion injection times for the survey scan and the MS/MS scans were 250 ms, respectively and the ion target values were set to 3e6 and 1e6, respectively. Selected sequenced ions were dynamically excluded for 60 sec.

### Data Search

Tandem mass spectra were extracted by Proteome Discoverer ver. 2.0. Charge state deconvolution and deisotoping were not performed. All MS/MS samples were analyzed using Mascot (Matrix Science, London, UK; version 2.4.1). Mascot was set up to search the Uniprot Human database (148,284 entries; downloaded on Oct. 9^th^, 2015) assuming the digestion enzyme trypsin. Mascot was searched with a fragment ion mass tolerance of 0.02 Da and a parent ion tolerance of 10 ppm. Methylthio of cysteine was specified in Mascot as a fixed modification. Phosphorylation of serine, threonine and tyrosine, polyamine sulfonamide (355.23 Da) of alanine, aspartic acid, glutamic acid, glycine, histidine, methionine, asparagine, glutamine, arginine, lysine, proline, valine, leucine, isoleucine, phenylalanine, tryptophan were specified in Mascot as variable modifications.

Scaffold (version Scaffold_4.2.1, Proteome Software Inc., Portland, OR) was used to validate MS/MS based peptide and protein identifications. Peptide identifications were accepted if they could be established at greater than 80.0% probability by the Peptide Prophet algorithm (Keller, A et al Anal. Chem. 2002;74(20):5383-92) with Scaffold delta-mass correction. Protein identifications were accepted if they could be established at greater than 95.0% probability and contained at least 1 identified peptide. Protein probabilities were assigned by the Protein Prophet algorithm (Nesvizhskii, Al et al Anal. Chem. 2003;75(17):4646-58). Proteins that contained similar peptides and could not be differentiated based on MS/MS analysis alone were grouped to satisfy the principles of parsimony. Proteins sharing significant peptide evidence were grouped into clusters.

### Compound Footprinting

Proteins were precipitated in 25mM ammonium bicarbonate, pH 8.0with Amicon Ultra-0.5 ml Centrifugal filter (EMD Millipore Inc., Billerica, MA, USA). For each sample, a total of 30 μg of protein were reduced with 40 mM DTT, alkylated with 100 mM of iodoacetamide, and trypsin-digested (at an enzyme to protein ratio (w/w) of 1:100). Tryptic digested peptides were desalted with C18-solid phase extraction. A hybrid quadrupole Orbitrap (Q Exactive plus) MS system (Thermo Fisher Scientific, Bremen, Germany) was used with high energy collision dissociation (HCD) in each MS and MS/MS cycle. The MS system was interfaced with an ultra-performance Easy-nLC 1000 system (Thermo Fisher Scientific, Bremen, Germany). 2 ug of sample was loaded onto a Acclaim Pepmap 100 pre-column (20 mm × 75 μm; 3 μm-C18) and then separated on a PepMap RSLC analytical column (500 mm × 75 μm; 2 μm-C18) at a flow rate of 300 nl/min of solvent A (0.1% formic acid), followed by a linear increase from 0% to 25% solvent B (0.1% formic acid, 99.9% acetonitrile) in 170 min and then ramping up to 98% B and stayed for 10 min. Then the system was equilibrated in solvent A for 30 min. The spectrum library was produced in the data dependent mode with survey scans acquired at a resolution of 70,000 at 200 m/z. The mass spectrometer was operated in MS/MS mode scanning from 350 to 1800 m/z. Up to the top 10 most abundant isotope patterns with charge 2 ∼ 5 from the survey scan were selected with an isolation window of 1.2 Th and fragmented by high energy collision dissociation (HCD) with normalized collision energies of 28. For directed LC–MS/MS, 14 inclusion mass lists comprising the ion masses of the observed precursor ions using Matrix-assisted laser desorption/ionization (MALDI) tandem time-of-flight (4700 Proteomic Analyzer, Applied Biosystems) were added (355.2300 m/z, 843.4582 m/z, 900.5524 m/z, 985.5220 m/z, 990.5589 m/z, 995.5890 m/z, 1114.6374 m/z, 1144.5905 m/z, 1171.5861 m/z, 1224.6189 m/z, 1358.7326 m/z, 1389.6503 m/z, 1488.7704 m/z, 1545.7766 m/z, 1572.8279 m/z, 1600.8084 m/z). The maximum ion injection times for the survey scan and the MS/MS scans were 250 ms, respectively and the ion target values were set to 3e6 and 1e6, respectively. Selected sequenced ions were dynamically excluded for 60 sec.

### Compound Footprinting Data Analysis

The raw MS/MS data files were processed by a thorough database searching considering biological modification and amino acid substitution against the Uniprot Human database (148,284 entries; downloaded on Oct. 9^th^, 2015) using Scaffod Q+S (Proteome Software Inc., Portland, OR, USA) and MASCOT 2.4 (Matrix Science Inc., Boston, MA, USA) with the following parameters: peptide tolerance at 10 ppm, tandem MS tolerance at ± 0.02 Da, peptide charge of 2+, 3+, 4+, trypsin as the enzyme, Carbamidomethyl (C) as fixed modifications, and Phosphorylation of serine, threonine and tyrosine, polyamine sulfonamide (355.23 Da) of alanine, aspartic acid, glutamic acid, glycine, histidine, methionine, asparagine, glutamine, arginine, lysine, proline, valine, leucine, isoleucine, phenylalanine, tryptophan were specified in Mascot as variable modifications. Peptide and protein were filtered using Scaffold Q+S with strict peptide and protein probabilities, 0.7 and 0.95, respectively.

### Surface-Plasmon Resonance Imaging (SPRi)

Recombinant HNRNPC, hnRNPA2/B1 and RALY were purchased from MyBiosource and Ray Biotech. Various concentrations of recombinant proteins dissolved in buffer were manually printed onto the bare gold-coated (thickness 47 nm) PlexArray Nanocapture Sensor Chip (Plexera Bioscience) at 40% humidity. Each concentration was printed in replicate, and each spot contained 0.2 μL of sample solution. The chip was incubated in 80% humidity at 4°C for overnight, and rinsed with 10×PBST for 10 min and deionized water two times for 10 min. The chip was then blocked with 5% (w/v) non-fat milk in water overnight, and washed with 10×PBST for 10 min, 1×PBST for 10 min, and deionized water twice for 10 min before being dried under a stream of nitrogen prior to use. SPRi measurements were performed with PlexAray HT (Plexera Bioscience). Collimated light (660 nm) is allowed to pass through the coupling prism, which then reflects off the SPR-active gold surface, and is received by the CCD camera. Buffers and samples were injected by a non-pulsatile piston pump into the 30 μL flow cell that was mounted on the coupling prism. Each measurement cycle contained four steps: washing with PBST running buffer at a constant rate of 2 μL/s to obtain a stable baseline, sample injection at 5 μL/s for binding, surface washing with PBST at 2 μL/s for 300 s, and regeneration with 0.5% (v/v) H3PO4 at 2 μL/s for 300 s. All the measurements were performed at 25°C. The signal changes after binding and washing (in AU) are recorded as the assay value. Selected protein-grafted regions in the SPR images were analyzed, and the average reflectivity variations of the chosen areas were plotted as a function of time. Real-time binding signals were recorded and analyzed by Data Analysis Module (DAM, Plexera Bioscience). Kinetic analysis was performed using BIAevaluation 4.1 software (Biacore, Inc.).

### Lentiviral Transduction of shRNA

Plasmids targeting HNRNPC were purchased from Dharmacon (GE Healthcare). A 2nd generation lentiviral packaging system was used to transfect 293T cells to produce viral particles. Briefly, 293T cells were grown to 70% confluency, after which the cells were transfected with shHNRNPC, or shScramble, along with psPAX2 and pMD2.G packaging and envelope virus using lipofecctamine 3000 (Invitrogen) in serum-free Opti-MEM media (Invitrogen). The viral supernatant was collected and used to transduce desired cell lines or aliquoted and stored at -80° C for future transductions. For effective transductions, the viral supernatant was supplemented with 8 µg/ml polybrene (Sigma) during both transductions. After 48 hours of transduction, puromycin selection was performed at concentration ranging between 0.5 µg/mL-2 µg/mL, depending on the cell line. Puromycin selection was performed for approximately 4-7 days, after which cells were used for further analysis.

### Growth Curve Analysis, Proliferation and Cell Viability analysis

To assess the proliferation and growth of cells transduced with shRNA, cells selected using puromycin for about 4-7 days were plated in a 6-well plate in selection media. Cells were counted using trypan blue exclusion method on alternative days for 10 days to establish the growth curve. Metabolic capacity of the cells was assessed using CellTiter Blue Assay (Promega) to examine the proliferation index of the cells. Cells were plated in a 96-well plate in 100 µL of their respective selection media. The cell titer reagent was added according to the manufacturer’s recommendations and fluorescence was measured using a microplate reader (BioTek Gen5) with excitation at 530-570 nm and emission at 580-620 nm. Cell viability was assessed using flow cytometric analysis using Annexin-V FITC and 7-AAD (BD Biosciences).

### Colony Forming Unit Assay

For primary specimens, 5,000 cells were re-suspended in 4 mL of Methocult media (STEMCELL Technologies). 1.1 mL of methocult-cell suspension was plated into a 6-well plate using 16-gauge blunt-end needles (STEMCELL Technologies). After 14 days, three types of colonies were quantified; Erythroid progenitor cells-colony forming unit-erythroid (CFU) and burst-forming unit-erythroid (BFU-E), Granulocyte and/or macrophage progenitor cells (CFU-granulocyte, macrophage (CFU-GM) and Multi-potential progenitor cells (CFU-granulocyte, erythrocyte, macrophage, megakaryocyte (CFU-GEMM). For cell lines, 10,000 cells were re-suspended in methocult media for subsequent plating and colonies were enumerated after 14 days.

### Proximity Ligation Assay

Proximity ligation assay (PLA) was performed using DuoLink PLA kit (Sigma) according to manufacturer’s instructions. Briefly, cells transduced with HNRNPC-wt or HNRNPC-kd were fixed using 100% methanol for 15 minutes at -20° C. The cells were washed, resuspended in 1X PBS and then smeared onto a glass slide. The cells were permeabilized using 0.3% Triton-X 100 for 10 minutes at room temperature. The cells were washed using 1% BSA in PBS and blocked using the blocking solution provided for 1 hour at 37° C. The slides were then stained with primary antibodies anti-c-Myc C33 antibody (SantaCruz) at 1:50 dilution and anti-Max antibody (Abcam) at 1:250 dilution for approximately 16 hours at 37° C. Following primary antibody incubation, the PLUS and MINUS probes (Rabbit and Mouse) were added, and subsequent steps were performed according to the manufacturer’s protocol. The slides were imaged using Nikon A1RMP confocal microscope (Nikon) and the images were quantified using ImageJ.

### Patient data

This study had the approval and guidance of the Institutional Review Boards (IRB) at Fox Chase Cancer Center, National Institutes of Health (NIH), Oregon Health & Science University (OHSU), Stanford University, University of Colorado, University of Florida, University of Kansas (KUMC), University of Miami, University of Texas Medical Center (UT Southwestern), and University of Utah. All patients gave informed consent to participate in this study. Data were processed as described in Tyner et al. 2018. TCGA data were obtained from the NIH National Cancer Institute GDC Legacy Archive. 1. The results published here are in whole or part based upon data generated by the Therapeutically Applicable Research to Generate Effective Treatments (https://www.cancer.gov/ccg/research/genome-sequencing/target)https: The data used for this analysis are available at the Genomic Data Commons (https://portal.gdc.cancer.gov).

### RNA Extraction and RNA Sequencing

RNA was extracted from cultured cells using Qiagen RNeasy mini kit as per the manufacturer’s protocol. Total RNA was quantified using NanoDrop spectrophotometer (Thermo Fisher). RNA sequencing was performed and analyzed by Novogene (novogene.com). DESeq2 was used to perform differential gene expression analysis. Genes with an adjusted P<0.05 and absolute log_2_ fold change>1 were described as differentially expressed genes.

### Read processing and quantification

Reads were paired-end 76bp and processed using FASTQC v0.11.7^32^ before trimming using Trimmomatic v0.36^33^ (LEADING:3, TRAILING:3, MINLEN:50, ILLUMINACLIP:TruSeq3-PE-2.fa:2:30:10:2:true, MAXINFO:50:0.3, SLIDINGWINDOW:4:15). Trimmed reads were subsequently mapped to the hg38 reference genome using STAR v2.6.1^34^ (--outFilterMultimapNmax 20, --outFilterMismatchNmax 999, --outFilterMismatchNoverReadLmax 0.04, --outSAMstrandField intronMotif, --alignIntronMin 20, --alignIntronMax 1000000, --alignMatesGapMax 1000000, --alignSJoverhangMin 8, --alignSJDBoverhangMin 1, --sjdbScore 1). Read quantification was performed using StringTie v1.3.4^35^ (-j 2) where independent assemblies were merged and re-quantified using the merged GTF.

### Transcript differential expression and alternative splicing

Count matrices were generated from both gene and transcript data within R v3.5.1 (R Core Team 2018) using tximport^36^. Differential gene and transcript quantification were performed using limma-voom^37^ (see R markdown for details) where features were filtered for low counts^38^, samples with abnormally low library sizes were removed and the remaining samples were normalized using TMM normalization. The results were filtered at FDR 0.05. Alternative splicing events were obtained using rMATS 4.0.2^39^and filtered at FDR 0.05.

### Data Analysis and Visualization

Data analysis was performed using Excel, CPython and R. Figures were generated using the CPython libraries seaborn and matplotlib and the R package limma. Sashimi plots were made using ggsashimi. Survival curves were made using the python library lifelines.

### Data availability

Read data were obtained through Synapse and are publicly available through the NIH/NCI dbGaP and Genomic Data Commons (GDC) resources. The dbGaP study ID is 30641 and accession ID is phs001657.v1.p1.

## SUPPLEMNTAL FIGURE LEGENDS

**Supplemental Figure 1.**
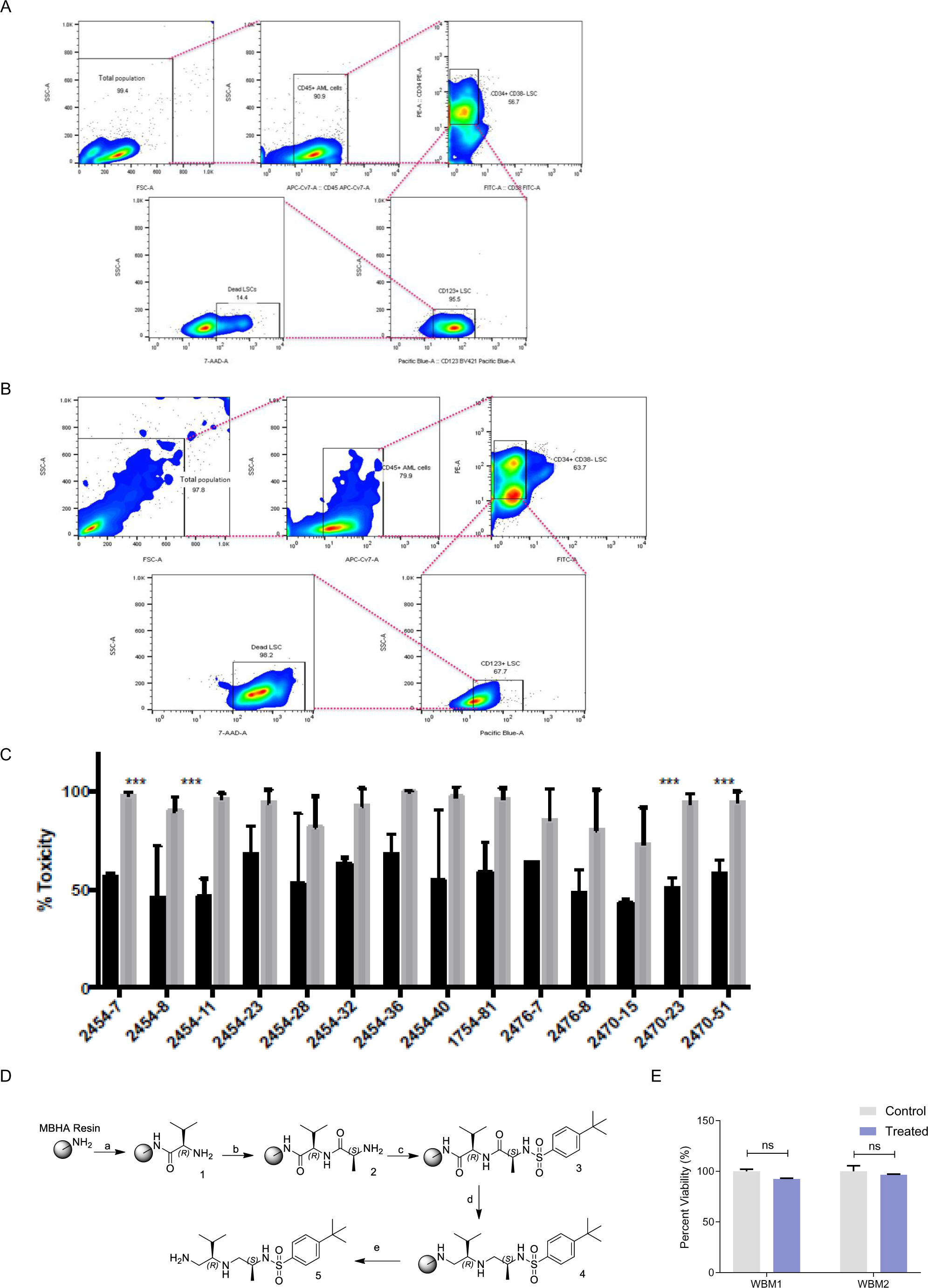
A-B) Gating strategy for analyzing leukemic stem cell toxicity using flow cytometry analysis. Total population of live cells were selected using forward (FSC) and side scatter (SSC). A subset of this population was identified as CD45+ AML cells, eliminating endothelial cell contamination. A subset of this population was sub-gated for CD34+, CD38-population, followed by further sub-gating to identify CD123+ leukemic stem cells. Live/dead analysis was performed using a combination of Annexin V and 7-Actinomycin A-D gating. C) Relative toxicity of top hits on leukemic stem cell population (LSC) in AML patient specimens and normal hematopoietic stem/ progenitor cells in human umbilical cord blood specimens (UCB). n=6 AML patient specimens and n=4 UCB specimens, error bars means ± s.e.m. *P<0.05, **P<0.01, ***P<0.001 and ****P<0.0001, Unpaired t-test. D) Synthesis of N-((S)-1-(((R)-1-amino-3-methylbutan-2-yl)amino)propan-2-yl)-4-(tert-butyl)benzenesulfonamide. (a) Boc-D-Val-OH (6 equiv), DIC (6 equiv), HOBt (6 equiv); 55%TFA/DCM; 5%DIE/DCM (b) Boc-L-Ala-OH (6 equiv), DIC (6 equiv), HOBt (6 equiv); 55%TFA/DCM; 5%DIEA/DCM (c) 4-tert-butylbenzene-1-sulfonyl chloride (10 equiv), DIEA (10 equiv) in 0.1M DCM; (d) BH3/THF (40 equiv), 65 °C, 72 h; piperidine, 65 °C, overnight; (e) anhydrous HF, 0 °C, 7 h. E) Percentage viability of mononuclear cells from normal adult bone marrow specimenS after 2470-51 treatment. n=2, error bars means ± s.e.m. *P<0.05, **P<0.01, ***P<0.001 and ****P<0.0001, Unpaired t-test.

**Supplemental Figure 2.**
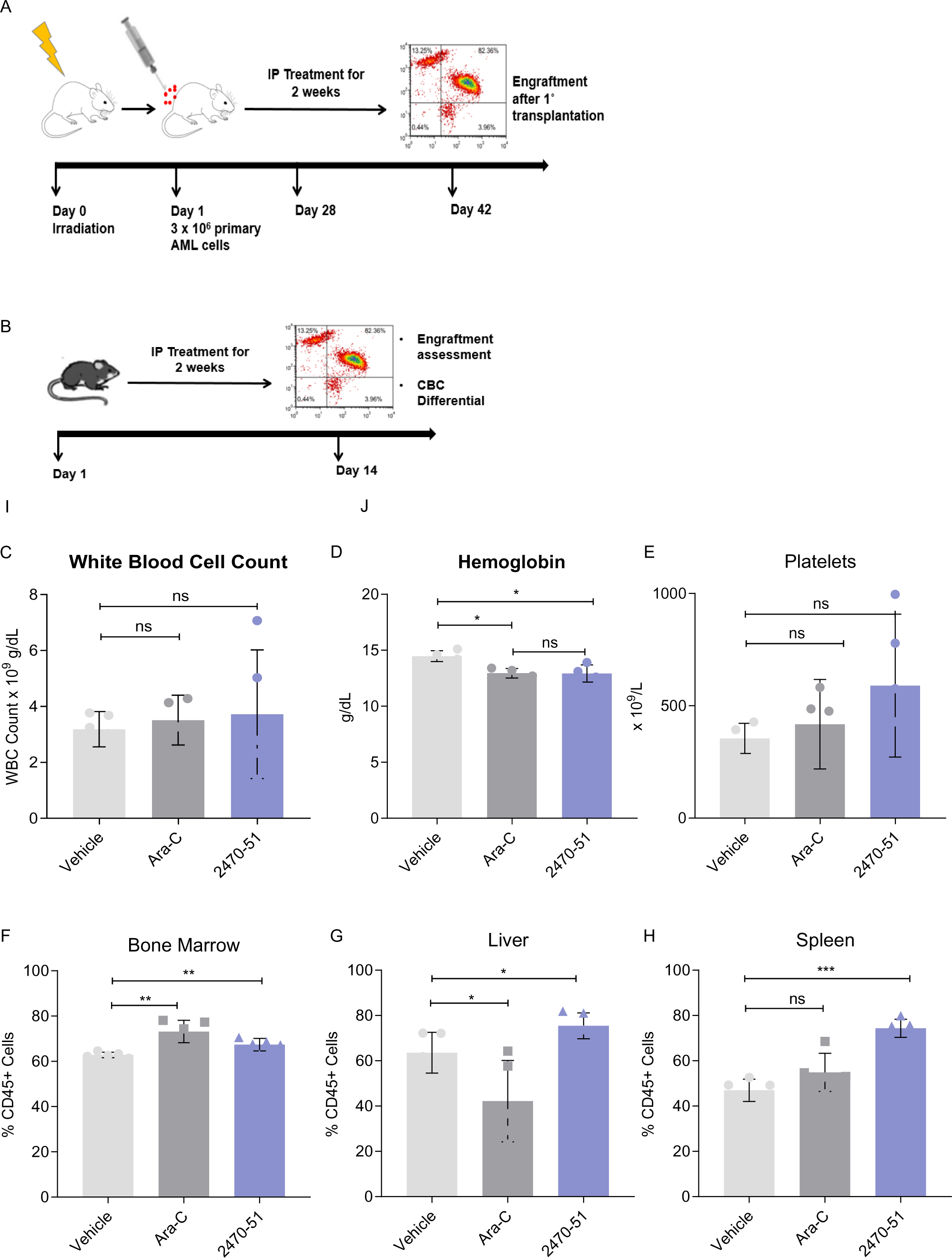
A) Schematic of *in vivo* treatment strategy to assess efficacy against AML. NSG mice were sub-lethally irradiated at 200cGy and transplanted with primary AML cells after 24 hours. After 4 weeks of allowing engraftment, mice were treated with Saline, Cytarabine (Ara-C) at 100 mg/kg or 2470-51 at 20 mg/kg using intra-peritoneal injections TIW for 2 weeks. Following 2 weeks of treatment, mice were euthanized, bone marrow, liver and spleens were harvested and assessed for AML engraftment using immunofluorescence (IF). n=8 mice per treatment group. B) Schematic of *in vivo treatment* strategy for assessing TPI2470-51 toxicity to normal cells. Mice were treated with Saline, Cytarabine (Ara-C) at 100 mg/kg or 2470-51 at 20 mg/kg using intra-peritoneal injections TIW for 2 weeks. Following 2 weeks of treatment, mice were euthanized, and peripheral blood was collected to quantify C) total white blood cell count, D) total hemoglobin, E) platelet levels. Bone marrow, liver and spleen were harvested to assess toxicity against normal human CD45+ cells (F-H).

**Supplemental Figure 3.**
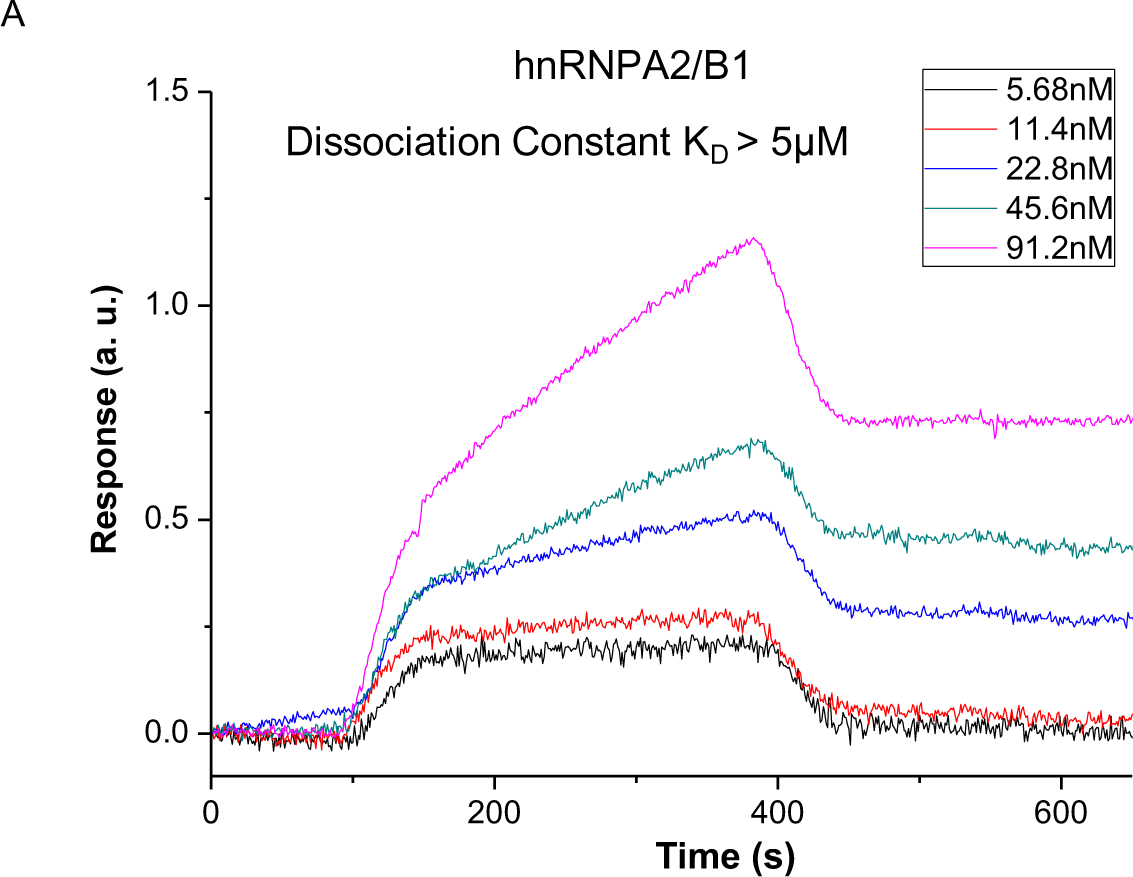
A) Surface plasmon resonance imaging (SPRi) to identify binding affinity between immobilized recombinant human hnRNPA2/B1 and 2470-51. Real-time binding signals were recorded and analyzed by Data Analysis Module (DAM). Kinetic analysis was performed using BIAevaluation 4.1 software. Association rate constant Ka, dissociation rate constant Kd and equilibrium dissociation constant KD values were calculated.

**Supplemental Figure 4.**
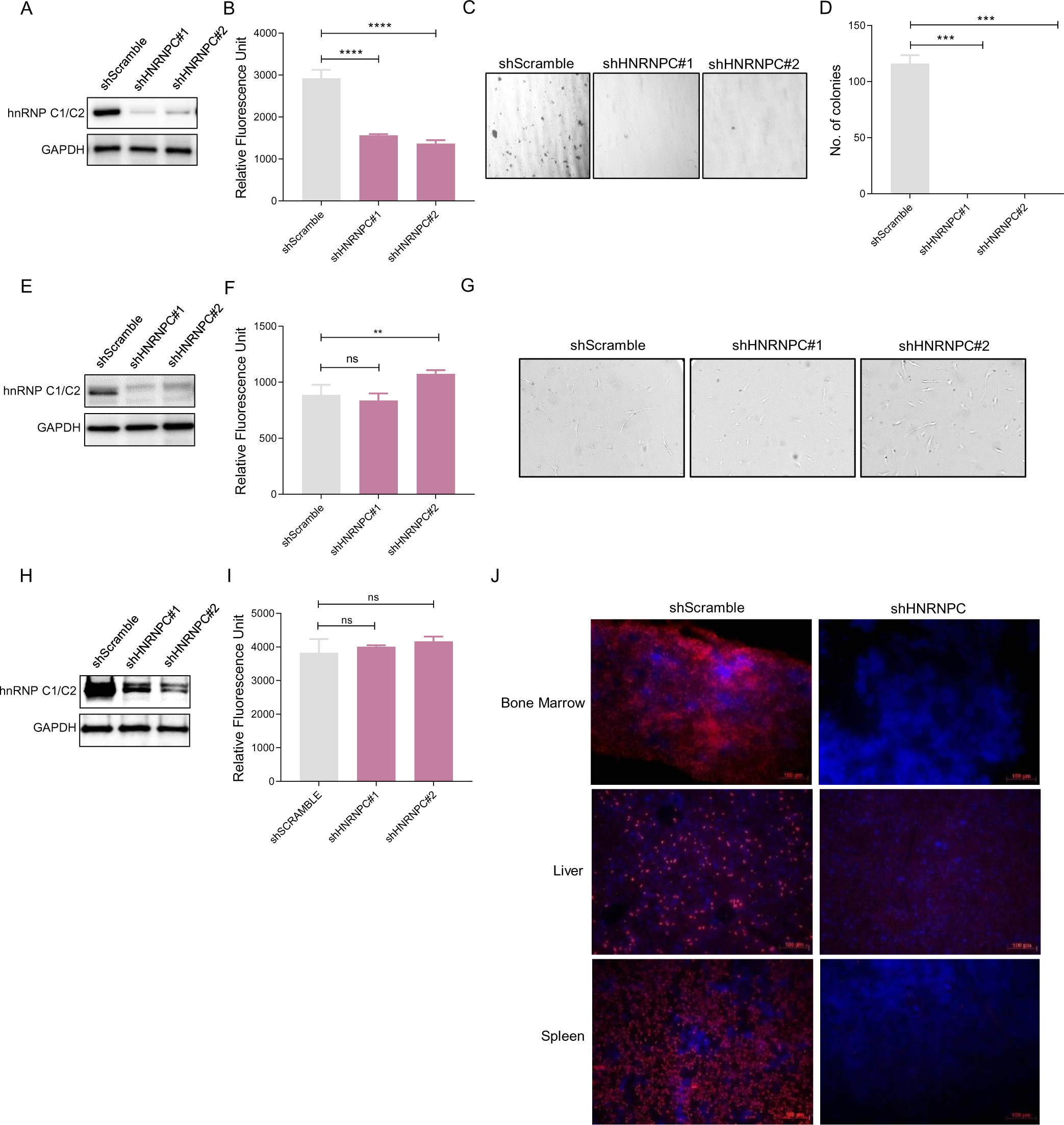
A) AML cell line THP-1 was transduced with lentiviruses expressing either a scrambled control (shScramble) or two independent shRNA targeting HNRNPC (shHNRNPC#1 and shHNRNPC#2) for 48 hours, followed by puromycin selection for 3-7 days. Puromycin resistant cells were immunoblotted for HNRNPC and Gapdh (loading control), day 5 after transduction. B) THP-1 cell metabolic activity using Cell titer blue assay as a surrogate for cell proliferation 7d post-transduction. Y-axis represents relative fluorescent units (RFU). n=3 independent experiments, error bars mean ± s.e.m. *P<0.05, **P<0.01, ***P<0.001 and ****P<0.0001, Unpaired t-test. C) Representative images of colony forming unit (CFU) assay of THP-1 cells 14d post-selection in a methylcellulose based medium. D) Number of colonies (>50 cells) counted 14d after plating in methylcellulose medium. Error bars means ± s.e.m. *P<0.05, **P<0.01, ***P<0.001 and ****P<0.0001, Unpaired t-test. E) Primary BMECs were transduced with lentiviruses expressing either a scrambled control (shScramble) or two independent shRNA targeting HNRNPC (shHNRNPC#1 and shHNRNPC#2) for 48 hours, followed by puromycin selection for 3-7 days. Puromycin resistant cells were immunoblotted for HNRNPC and Gapdh (loading control), day 5 after transduction. F) BMEC cell metabolic activity using Cell titer blue assay as a surrogate for cell proliferation 7d post-transduction. Y-axis represents relative fluorescent units (RFU). n=3 independent experiments, error bars mean ± s.e.m. *P<0.05, **P<0.01, ***P<0.001 and ****P<0.0001, Unpaired t-test. G) Representative phase-contrast images of BMEC cells 7d post transduction. H) Normal fibroblast-like cell line HS-5 was transduced with lentiviruses expressing either a scrambled control (shScramble) or two independent shRNA targeting HNRNPC (shHNRNPC#1 and shHNRNPC#2) for 48 hours, followed by puromycin selection for 3-7 days. Puromycin resistant cells were immunoblotted for HNRNPC and Gapdh (loading control), day 5 after transduction. I) HS-5 cell metabolic activity using Cell titer blue assay as a surrogate for cell proliferation 7d post-transduction. Y-axis represents relative fluorescent units (RFU). n=3 independent experiments, error bars mean ± s.e.m. *P<0.05, **P<0.01, ***P<0.001 and ****P<0.0001, Unpaired t-test. J) Representative IF images of bone marrow, liver and spleen harvested from NSG mice engrafted with MV4-11 AML cell line transduced with shScramble or shHNRNPC lentivirus. Blue=DAPI, Red=hCD45+ AML cells. n=6 mice per cohort.

**Supplemental Figure 5.**
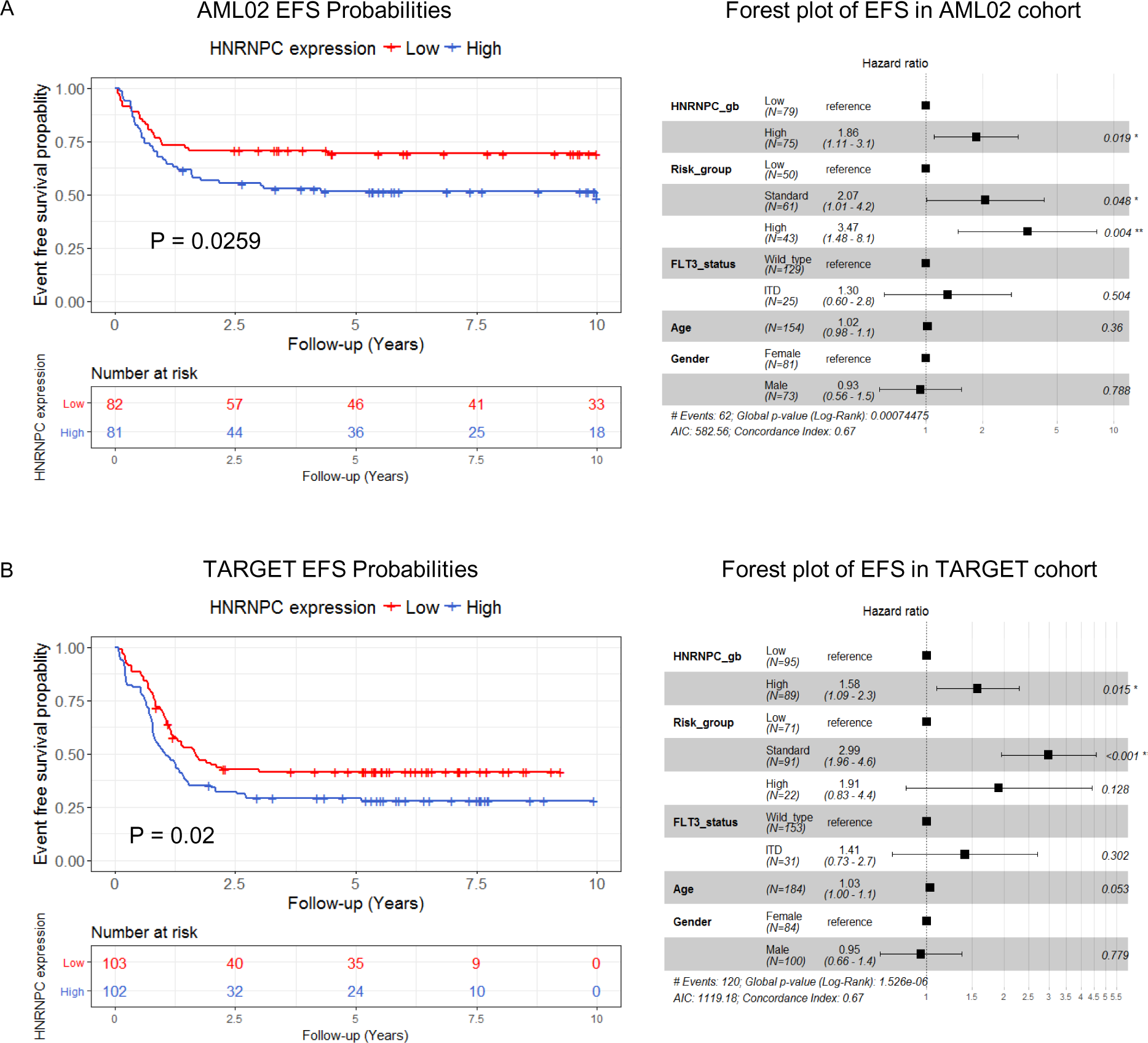
Survival probabilities of A) AML-02 and B) TARGET-AML cohort based on HNRNPC expression. EFS probabilities of patients based on HNRNPC expression. Forest plot indicating the hazard ratios and P-values for each variable used in the multi-variate survival analysis.

**Supplemental Figure 6.**
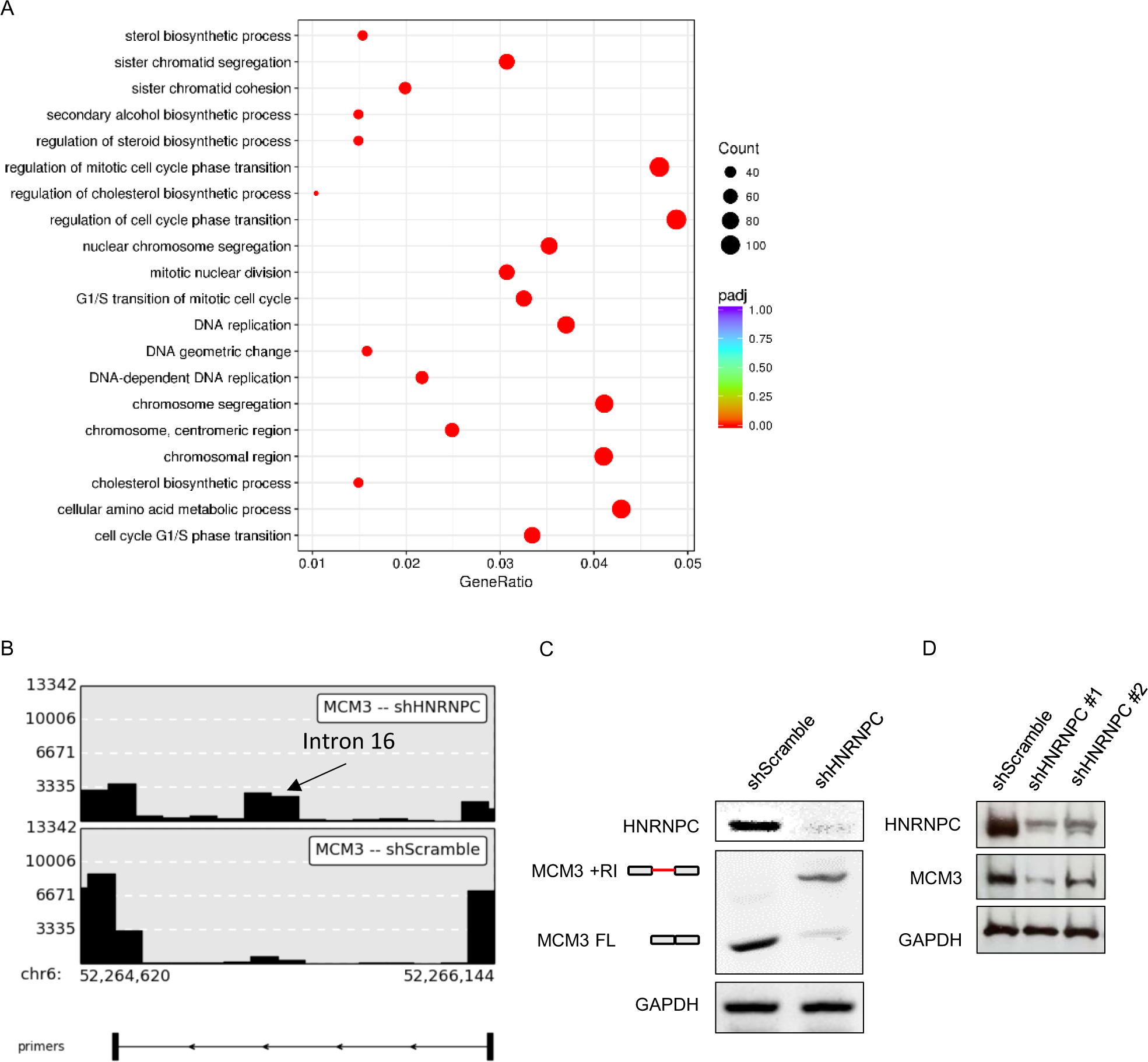
A) Functional enrichment analysis of genes downregulated after HNRNPC depletion. Dot plot indicates the enrichment of genes to pathways using Gene Ontology analysis. B) RT-PCR validation of intron 16 retention event in MCM3. PrimerSeq software was used to design primers in exons flanking intron 16. C) RT-PCR of MCM3 along with HNRNPC and *GAPDH* levels included for reference. D) Western blotting to visualize MCM3 protein expression after HNRNPC depletion.

**Supplemental Figure 7.**
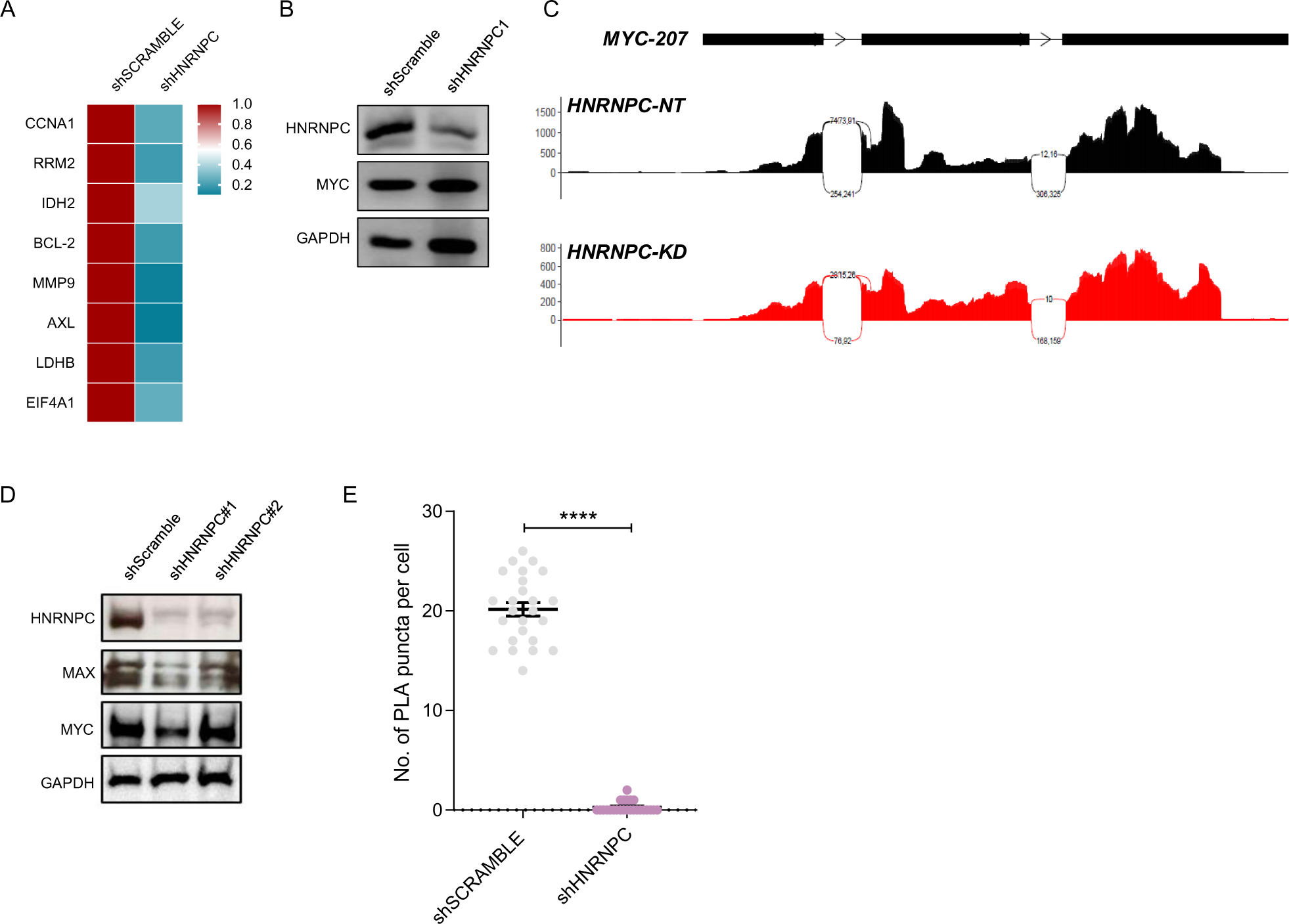
A) Heatmap showing expression of MYC targets after HNRNPC depletion. Heatmap indicates calculated z-scores. B) mRNA levels of *MYC*, HNRNPC and *GAPDH* analyzed using RT-PCR. C) Sashimi plot representing the splicing pattern of *MYC* in shScramble (Black) and shHNRNPC (Red) THP-1 cells. D) Western blotting of MAX and MYC protein levels after HNRNPC depletion. E) Quantification of PLA puncta per cell. n=25 cells per condition.

**Supplemental Figure 8.**
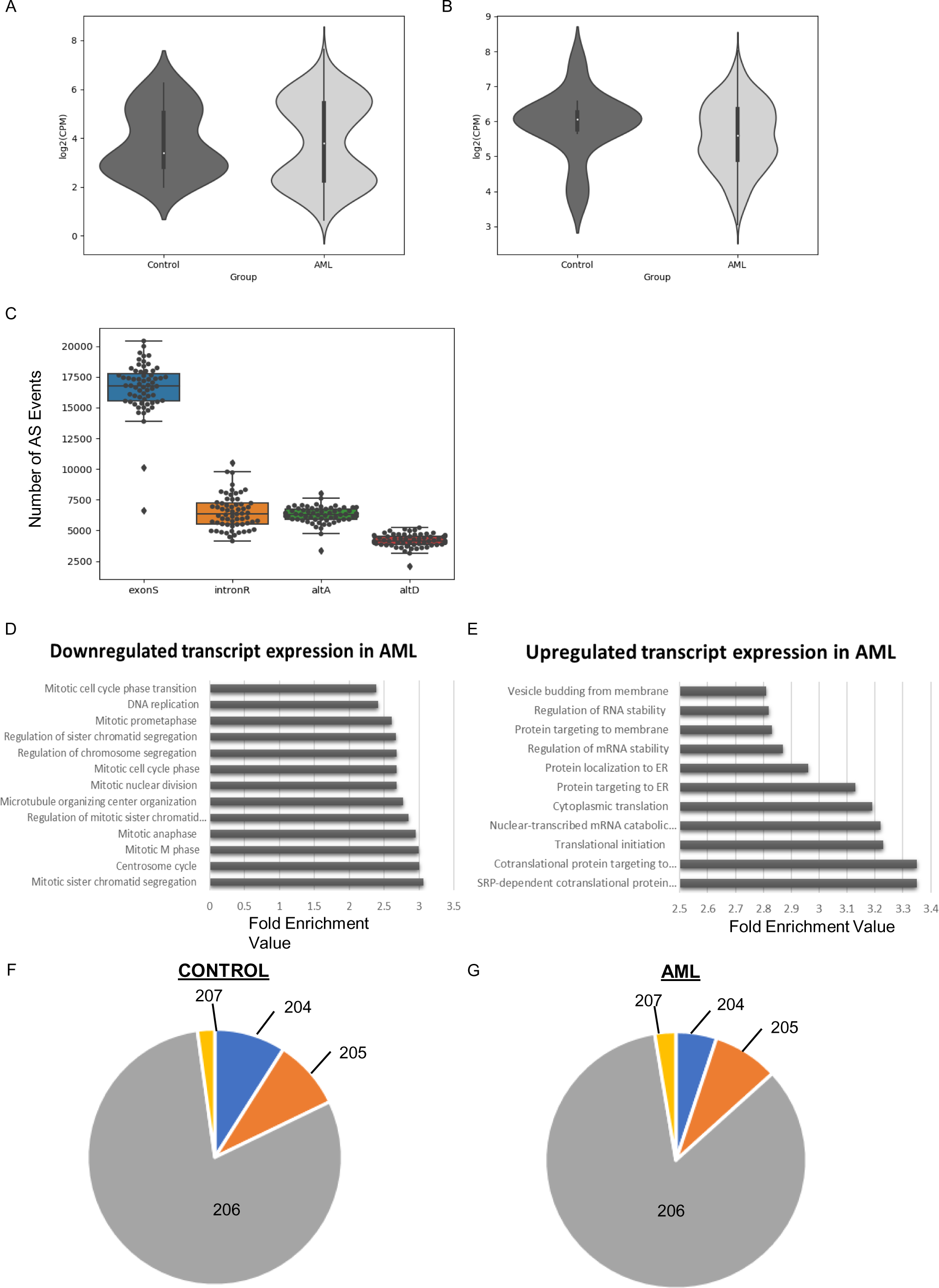
A) MAX and B) *MYC* mRNA levels in control and AML patients in the BEAT-AML patient dataset. C) Quantification of alternative splicing events comparing control and AML patients analyzed using RMATS. exonS=Skipped Exons, intronR= Retained Introns, altA= Alternative 5’ acceptor site, altD= Alternative 5’ donor site. D-E) Functional enrichment analysis of differentially expressed transcripts comparing normal and AML patients. F-G) Proportional distribution of *MYC* transcripts in control and AML patients. Each segment represents a distinct *MYC* isoform.

## REFERENCES

1. Döhner, H. et al. Diagnosis and management of AML in adults: 2017 ELN recommendations from an international expert panel. Blood 129, 424–447, doi:10.1182/blood-2016-08-733196 (2017).

2. Cogle, C. R. et al. Functional integration of acute myeloid leukemia into the vascular niche. Leukemia, doi:10.1038/leu.2014.109 (2014).

3. Cogle CR, Bosse RC, Brewer T, Migdady Y, Shirzad R, Kampen KR, Saki N. Acute myeloid leukemia in the vascular niche. Cancer Lett. 2016 Oct 1;380(2):552–560. doi: 10.1016/j.canlet.2015.05.007. Epub 2015 May 8. PMID: 25963886.

4. Duarte D, Hawkins ED, Lo Celso C. The interplay of leukemia cells and the bone marrow microenvironment. Blood. 2018 Apr 5;131(14):1507–1511. doi: 10.1182/blood-2017-12-784132. Epub 2018 Feb 27. PMID: 29487069.

5. Winkler IG, Barbier V, Nowlan B, Jacobsen RN, Forristal CE, Patton JT, Magnani JL, Lévesque JP. Vascular niche E-selectin regulates hematopoietic stem cell dormancy, self renewal and chemoresistance. Nat Med. 2012 Nov;18(11):1651–7. doi: 10.1038/nm.2969. Epub 2012 Oct 21. PMID: 23086476.

6. Ross PL, Huang YN, Marchese JN, Williamson B, Parker K, Hattan S, Khainovski N, Pillai S, Dey S, Daniels S, Purkayastha S, Juhasz P, Martin S, Bartlet-Jones M, He F, Jacobson A, Pappin DJ. Multiplexed protein quantitation in Saccharomyces cerevisiae using amine-reactive isobaric tagging reagents. Mol Cell Proteomics. 2004 Dec;3(12):1154–69. doi: 10.1074/mcp.M400129-MCP200. Epub 2004 Sep 22. PMID: 15385600.

7. Pai MY, Lomenick B, Hwang H, Schiestl R, McBride W, Loo JA, Huang J. Drug affinity responsive target stability (DARTS) for small-molecule target identification. Methods Mol Biol. 2015;1263:287–98. doi: 10.1007/978-1-4939-2269-7_22. PMID: 25618353; PMCID: PMC4442491.

8. Lomenick B, Jung G, Wohlschlegel JA, Huang J. Target identification using drug affinity responsive target stability (DARTS). Curr Protoc Chem Biol. 2011 Dec 1;3(4):163–180. doi: 10.1002/9780470559277.ch110180. PMID: 22229126; PMCID: PMC3251962.

9. Lomenick B, Hao R, Jonai N, Chin RM, Aghajan M, Warburton S, Wang J, Wu RP, Gomez F, Loo JA, Wohlschlegel JA, Vondriska TM, Pelletier J, Herschman HR, Clardy J, Clarke CF, Huang J. Target identification using drug affinity responsive target stability (DARTS). Proc Natl Acad Sci U S A. 2009 Dec 22;106(51):21984–9. doi: 10.1073/pnas.0910040106. Epub 2009 Dec 7. PMID: 19995983; PMCID: PMC2789755.

10. Choi YD, Grabowski PJ, Sharp PA, Dreyfuss G. Heterogeneous nuclear ribonucleoproteins: role in RNA splicing. Science. 1986 Mar 28;231(4745):1534-9. doi: 10.1126/science.3952495. PMID: 3952495.

11. Zarnack K, König J, Tajnik M, Martincorena I, Eustermann S, Stévant I, Reyes A, Anders S, Luscombe NM, Ule J. Direct competition between hnRNP C and U2AF65 protects the transcriptome from the exonization of Alu elements. Cell. 2013 Jan 31;152(3):453–66. doi: 10.1016/j.cell.2012.12.023. PMID: 23374342; PMCID: PMC3629564.

12. Holcík M, Gordon BW, Korneluk RG. The internal ribosome entry site-mediated translation of antiapoptotic protein XIAP is modulated by the heterogeneous nuclear ribonucleoproteins C1 and C2. Mol Cell Biol. 2003 Jan;23(1):280–8. doi: 10.1128/MCB.23.1.280-288.2003. PMID: 12482981; PMCID: PMC140676.

13. Schepens B, Tinton SA, Bruynooghe Y, Parthoens E, Haegman M, Beyaert R, Cornelis S. A role for hnRNP C1/C2 and Unr in internal initiation of translation during mitosis. EMBO J. 2007 Jan 10;26(1):158–69. doi: 10.1038/sj.emboj.7601468. Epub 2006 Dec 7. PMID: 17159903; PMCID: PMC1782369.

14. Kim JH, Paek KY, Choi K, Kim TD, Hahm B, Kim KT, Jang SK. Heterogeneous nuclear ribonucleoprotein C modulates translation of c-myc mRNA in a cell cycle phase-dependent manner. Mol Cell Biol. 2003 Jan;23(2):708–20. doi: 10.1128/MCB.23.2.708-720.2003. PMID: 12509468; PMCID: PMC151538.

15. Navickas A, Asgharian H, Winkler J, Fish L, Garcia K, Markett D, Dodel M, Culbertson B, Miglani S, Joshi T, Yin K, Nguyen P, Zhang S, Stevers N, Hwang HW, Mardakheh F, Goga A, Goodarzi H. An mRNA processing pathway suppresses metastasis by governing translational control from the nucleus. Nat Cell Biol. 2023 Jun;25(6):892–903. doi: 10.1038/s41556-023-01141-9. Epub 2023 May 8. PMID: 37156909; PMCID: PMC10264242. Michaud EJ, Bultman SJ, Stubbs LJ, Woychik RP. The embryonic lethality of homozygous lethal yellow mice (Ay/Ay) is associated with the disruption of a novel RNA-binding protein. Genes Dev. 1993 Jul;7(7A):1203–13. doi: 10.1101/gad.7.7a.1203. PMID: 8319910.

16. Duhl DM, Stevens ME, Vrieling H, Saxon PJ, Miller MW, Epstein CJ, Barsh GS. Pleiotropic effects of the mouse lethal yellow (Ay) mutation explained by deletion of a maternally expressed gene and the simultaneous production of agouti fusion RNAs. Development. 1994 Jun;120(6):1695–708. doi: 10.1242/dev.120.6.1695. PMID: 8050375.

17. Williamson DJ, Banik-Maiti S, DeGregori J, Ruley HE. hnRNP C is required for postimplantation mouse development but Is dispensable for cell viability. Mol Cell Biol. 2000 Jun;20(11):4094–105. doi: 10.1128/MCB.20.11.4094-4105.2000. PMID: 10805751; PMCID: PMC85779.

18. Niggl E, Bouman A, Briere LC, Hoogenboezem RM, Wallaard I, Park J, Admard J, Wilke M, Harris-Mostert EDRO, Elgersma M, Bain J, Balasubramanian M, Banka S, Benke PJ, Bertrand M, Blesson AE, Clayton-Smith J, Ellingford JM, Gillentine MA, Goodloe DH, Haack TB, Jain M, Krantz I, Luu SM, McPheron M, Muss CL, Raible SE, Robin NH, Spiller M, Starling S, Sweetser DA, Thiffault I, Vetrini F, Witt D, Woods E, Zhou D; Genomics England Research Consortium; Undiagnosed Diseases Network; Elgersma Y, van Esbroeck ACM. HNRNPC haploinsufficiency affects alternative splicing of intellectual disability-associated genes and causes a neurodevelopmental disorder. Am J Hum Genet. 2023 Aug 3;110(8):1414–1435. doi: 10.1016/j.ajhg.2023.07.005. PMID: 37541189; PMCID: PMC10432175.

19. Cancer Genome Atlas Research Network; Ley TJ, Miller C, Ding L, Raphael BJ, Mungall AJ, Robertson A, Hoadley K, Triche TJ Jr, Laird PW, Baty JD, Fulton LL, Fulton R, Heath SE, Kalicki-Veizer J, Kandoth C, Klco JM, Koboldt DC, Kanchi KL, Kulkarni S, Lamprecht TL, Larson DE, Lin L, Lu C, McLellan MD, McMichael JF, Payton J, Schmidt H, Spencer DH, Tomasson MH, Wallis JW, Wartman LD, Watson MA, Welch J, Wendl MC, Ally A, Balasundaram M, Birol I, Butterfield Y, Chiu R, Chu A, Chuah E, Chun HJ, Corbett R, Dhalla N, Guin R, He A, Hirst C, Hirst M, Holt RA, Jones S, Karsan A, Lee D, Li HI, Marra MA, Mayo M, Moore RA, Mungall K, Parker J, Pleasance E, Plettner P, Schein J, Stoll D, Swanson L, Tam A, Thiessen N, Varhol R, Wye N, Zhao Y, Gabriel S, Getz G, Sougnez C, Zou L, Leiserson MD, Vandin F, Wu HT, Applebaum F, Baylin SB, Akbani R, Broom BM, Chen K, Motter TC, Nguyen K, Weinstein JN, Zhang N, Ferguson ML, Adams C, Black A, Bowen J, Gastier-Foster J, Grossman T, Lichtenberg T, Wise L, Davidsen T, Demchok JA, Shaw KR, Sheth M, Sofia HJ, Yang L, Downing JR, Eley G. Genomic and epigenomic landscapes of adult de novo acute myeloid leukemia. N Engl J Med. 2013 May 30;368(22):2059–74. doi: 10.1056/NEJMoa1301689. Epub 2013 May 1. Erratum in: N Engl J Med. 2013 Jul 4;369(1):98. PMID: 23634996; PMCID: PMC3767041.

20. König J, Zarnack K, Rot G, Curk T, Kayikci M, Zupan B, Turner DJ, Luscombe NM, Ule J. iCLIP reveals the function of hnRNP particles in splicing at individual nucleotide resolution. Nat Struct Mol Biol. 2010 Jul;17(7):909–15. doi: 10.1038/nsmb.1838. Epub 2010 Jul 4. PMID: 20601959; PMCID: PMC3000544.

21. 21. Jan Attig, Igor Ruiz de los Mozos, Nejc Haberman, Zhen Wang, Warren Emmett, Kathi Zarnack, Julian König, Jernej Ule (2016) Splicing repression allows the gradual emergence of new Alu-exons in primate evolution eLife 5:e19545

22. Walhout AJ, Gubbels JM, Bernards R, van der Vliet PC, Timmers HT. c-Myc/Max heterodimers bind cooperatively to the E-box sequences located in the first intron of the rat ornithine decarboxylase (ODC) gene. Nucleic Acids Res. 1997 Apr 15;25(8):1493–501. doi: 10.1093/nar/25.8.1493. PMID: 9162900; PMCID: PMC146624.

23. Kato GJ, Lee WM, Chen LL, Dang CV. Max: functional domains and interaction with c-Myc. Genes Dev. 1992 Jan;6(1):81–92. doi: 10.1101/gad.6.1.81. PMID: 1730412.

24. Dang CV, McGuire M, Buckmire M, Lee WM. Involvement of the ’leucine zipper’ region in the oligomerization and transforming activity of human c-myc protein. Nature. 1989 Feb 16;337(6208):664-6. doi: 10.1038/337664a0. PMID: 2645525.

25. Hoffman B, Amanullah A, Shafarenko M, Liebermann DA. The proto-oncogene c-myc in hematopoietic development and leukemogenesis. Oncogene. 2002 May 13;21(21):3414–21. doi: 10.1038/sj.onc.1205400. PMID: 12032779.

26. Luo H, Li Q, O’Neal J, Kreisel F, Le Beau MM, Tomasson MH. c-Myc rapidly induces acute myeloid leukemia in mice without evidence of lymphoma-associated antiapoptotic mutations. Blood. 2005 Oct 1;106(7):2452–61. doi: 10.1182/blood-2005-02-0734. Epub 2005 Jun 21. PMID: 15972450.

27. Chen J, Odenike O, Rowley JD. Leukaemogenesis: more than mutant genes. Nat Rev Cancer. 2010 Jan;10(1):23–36. doi: 10.1038/nrc2765. PMID: 20029422; PMCID: PMC2972637.

28. Tyner JW, Tognon CE, Bottomly D, Wilmot B, Kurtz SE, Savage SL, Long N, Schultz AR, Traer E, Abel M, Agarwal A, Blucher A, Borate U, Bryant J, Burke R, Carlos A, Carpenter R, Carroll J, Chang BH, Coblentz C, d’Almeida A, Cook R, Danilov A, Dao KT, Degnin M, Devine D, Dibb J, Edwards DK 5th, Eide CA, English I, Glover J, Henson R, Ho H, Jemal A, Johnson K, Johnson R, Junio B, Kaempf A, Leonard J, Lin C, Liu SQ, Lo P, Loriaux MM, Luty S, Macey T, MacManiman J, Martinez J, Mori M, Nelson D, Nichols C, Peters J, Ramsdill J, Rofelty A, Schuff R, Searles R, Segerdell E, Smith RL, Spurgeon SE, Sweeney T, Thapa A, Visser C, Wagner J, Watanabe-Smith K, Werth K, Wolf J, White L, Yates A, Zhang H, Cogle CR, Collins RH, Connolly DC, Deininger MW, Drusbosky L, Hourigan CS, Jordan CT, Kropf P, Lin TL, Martinez ME, Medeiros BC, Pallapati RR, Pollyea DA, Swords RT, Watts JM, Weir SJ, Wiest DL, Winters RM, McWeeney SK, Druker BJ. Functional genomic landscape of acute myeloid leukaemia. Nature. 2018 Oct;562(7728):526–531. doi: 10.1038/s41586-018-0623-z. Epub 2018 Oct 17. PMID: 30333627; PMCID: PMC6280667.

29. Debevec G, Chen W, Yu Y, Houghten RA, Giulianotti MA. Libraries from Libraries: A Series of Sulfonamide Linked Heterocycles Derived from the Same Scaffold. Tetrahedron Lett. 2013 Aug 7;54(32):10.1016/j.tetlet.2013.06.003. doi: 10.1016/j.tetlet.2013.06.003. PMID: 24363466; PMCID: PMC3867824.

30. Pinilla C, Appel JR, Borràs E, Houghten RA. Advances in the use of synthetic combinatorial chemistry: mixture-based libraries. Nat Med. 2003 Jan;9(1):118–22. doi: 10.1038/nm0103-118. PMID: 12514724.

31. Houghten, R.A., General method for the rapid solid-phase synthesis of large numbers of peptides: specificity of antigen-antibody interaction at the level of individual amino acids. Proceedings of the National Academy of Sciences, 1985. 82(15): p. 5131.

32. 32. Andrews S. (2010). FastQC: a quality control tool for high throughput sequence data. Available online at: http://www.bioinformatics.babraham.ac.uk/projects/fastqc

33. Bolger AM, Lohse M, Usadel B. Trimmomatic: a flexible trimmer for Illumina sequence data. Bioinformatics. 2014 Aug 1;30(15):2114–20. doi: 10.1093/bioinformatics/btu170. Epub 2014 Apr 1. PMID: 24695404; PMCID: PMC4103590.

34. Dobin A, Davis CA, Schlesinger F, Drenkow J, Zaleski C, Jha S, Batut P, Chaisson M, Gingeras TR. STAR: ultrafast universal RNA-seq aligner. Bioinformatics. 2013 Jan 1;29(1):15–21. doi: 10.1093/bioinformatics/bts635. Epub 2012 Oct 25. PMID: 23104886; PMCID: PMC3530905.

35. Pertea M, Pertea GM, Antonescu CM, Chang TC, Mendell JT, Salzberg SL. StringTie enables improved reconstruction of a transcriptome from RNA-seq reads. Nat Biotechnol. 2015 Mar;33(3):290–5. doi: 10.1038/nbt.3122. Epub 2015 Feb 18. PMID: 25690850; PMCID: PMC4643835.

36. Soneson C, Love MI, Robinson MD. Differential analyses for RNA-seq: transcript-level estimates improve gene-level inferences. F1000Res. 2015 Dec 30;4:1521. doi: 10.12688/f1000research.7563.2. PMID: 26925227; PMCID: PMC4712774.

37. 37. Law, C.W., Chen, Y., Shi, W., et al. voom: precision weights unlock linear model analysis tools for RNA-seq read counts. Genome Biol 15, R29 (2014). 10.1186/gb-2014-15-2-r29

38. 38. Robinson, M.D., Oshlack, A. A scaling normalization method for differential expression analysis of RNA-seq data. Genome Biol 11, R25 (2010). 10.1186/gb-2010-11-3-r25

39. Shen S, Park JW, Lu ZX, Lin L, Henry MD, Wu YN, Zhou Q, Xing Y. rMATS: robust and flexible detection of differential alternative splicing from replicate RNA-Seq data. Proc Natl Acad Sci U S A. 2014 Dec 23;111(51):E5593–601. doi: 10.1073/pnas.1419161111. Epub 2014 Dec 5. PMID: 25480548; PMCID: PMC4280593.

40. Figures 1A, 2A, 2E, 3I and Supplementary Figures 2A and 2B were created using Biorender.com

